# Contrasting patterns of single nucleotide polymorphisms and structural variations across multiple invasions

**DOI:** 10.1101/2022.07.04.498653

**Authors:** Katarina C. Stuart, Richard J. Edwards, William B. Sherwin, Lee A. Rollins

## Abstract

Adaptive divergence is a fundamental process that shapes genetic diversity within and across species. Structural variants (SVs) are large-scale genetic differences (insertion, deletions, and rearrangements) within a species or population. SVs can cause important functional differences in the individual’s phenotype. Characterising SVs across invasive species will help fill knowledge gaps regarding how patterns of genetic diversity and genetic architecture shape rapid adaptation in response to new selection regimes. In this project we seek to understand patterns in genetic diversity within the globally invasive European starling, *Sturnus vulgaris*. We use whole genome sequencing of eight native United Kingdom (UK), eight invasive North America (NA), and 33 invasive Australian (AU) starlings to examine patterns in genome-wide SNPs and SVs between populations and within Australia. The findings of our research demonstrate that even within recently diverged lineages or populations, there may be high amounts of structural variation. Further, patterns of genetic diversity estimated from SVs do not necessarily reflect relative patterns from SNP data, either when considering patterns of diversity along the length of the organism’s chromosomes (owing to enrichment of SVs in sub telomeric repeat regions), or interpopulation diversity patterns (possibly a result of altered selection regimes or introduction history). Finally, we find that levels of balancing selection within the native range differ across SNP and SV of different classes and outlier classifications. Overall, our results demonstrate that the processes that shape allelic diversity within populations is complex and supports the need for further investigation of SVs across a range of taxa to better understand correlations between oft well studied SNP diversity and that of SVs.

## 1. Introduction

Adaptive divergence is a fundamental process that shapes genetic diversity within and across species (Schluter 2000; Arnold *et al*. 2001). Much remains unknown about the proximate mechanisms that underlie adaptive divergence, and the interacting roles of genetic diversity, gene flow, and selection in determining evolutionary trajectories (Campbell *et al*. 2018). Further, these evolutionary changes occasionally occur over very short biological timescales, a phenomenon termed rapid adaptation or rapid evolution (Thompson 1998). Central to this are questions about how genetic variants arise and remain within a population without being deleterious to individual fitness (Agrawal & Whitlock 2012), and conversely how these different genetic variants may respond to selective regimes (Kimura 1991; Eyre-Walker & Keightley 2007). Changes in the frequencies of genetic variants may result from balancing selection, which enables adaptation to an intermediate or changeable environment, or from directional selection, which enables adaptation to an environment that is locally constant (Hedrick 2007). Concurrently, there is a growing appreciation for the potential functional role of the non-protein coding regions that make up 98% of most eukaryotic genomes (Shihab *et al*. 2015), including non-coding regions that have important functional impacts on an organism’s phenotype, as shown in many studies (Zhang *et al*. 2018).

One major class of genetic mutations is structural variants (SVs). SVs are large-scale mutations within an organism’s genome that can result in important functional differences in the individual’s phenotype. Due to their size, single SVs have more potential for multiple impacts than single nucleotide variants (SNPs), and often encompass more of the genome (Alkan *et al*. 2011; Alonge *et al*. 2020). SVs encompass a wide variety of different genetic variants, including deletions (DEL), insertions (INS), duplications (DUP), inversions (INV), and translocations (TRA) (Alkan *et al*. 2011; Alonge *et al*. 2020). SVs may have a profound impact on the organisms’ phenotype, through direct alteration of gene content (Zhao *et al*. 2020) or alteration of transcription levels of adjacent genes (Puig *et al*. 2004). Examples of dramatic SV-affected phenotypes include altered pathogen resistance (Hämälä *et al*. 2021), modified plant flowering times (Todesco *et al*. 2020), and reproductive strategies (Lamichhaney *et al*. 2016).

The study of SVs has been somewhat limited until recent years, both due to their previously supposed rarity as well as the computational difficulties in characterising them (Ho *et al*. 2020). A range of bioinformatic process now exist through which people may characterise structural variants across a variety of sequencing data types including short-read, long-read, and RNA-seq (Mahmoud *et al*. 2019; Ho *et al*. 2020). These technical developments are allowing the role SVs play in adaptive divergence and speciation to be examined, and also provide an opportunity to contrast neutral and adaptive patterns in population genetic structure and diversity using SNP and SV data across a wide range of taxa. Within humans, SNPs explain less than a tenth of the between-individual genetic variation when considering all genetic variants, including SVs (Pang *et al*. 2010). Within the marine teleost *Chrysophrys auratus*, SV coverage of the genome was threefold that of SNPs (Catanach *et al*. 2019), and within chocolate trees SV coverage is eight times that of SNPs (Hämälä *et al*. 2021). While absolute comparisons of relative SV and SNP numbers across studies may be difficult due to different sequencing and data processing approaches, it is likely that SVs may play an important role in adaptation, with their effects being unable to be captured by SNP analysis alone (Dorant *et al*. 2020). Understanding the adaptive role SVs play in populations is hence of vital importance for understand standing genetic variation and the adaptive capacity of a population.

The importance of understanding patterns in SVs extends to invasive species, where they serve as a potential mechanism for rapid evolution that is often seen in invasive ranges (Colautti & Barrett 2013; Vandepitte *et al*. 2014; Rollins *et al*. 2015). It is even possible for the translocation and invasion process itself, by causing physiological stress in response to novel environments, to induce SV movement (Stapley *et al*. 2015). Characterising SVs across invasive species will help fill in knowledge gaps regarding patterns of genetic diversity and genetic architecture within successful invasive populations (Bock *et al*. 2015). Studies of SVs that try to explain a phenotype difference between organisms (e.g. finding a large explanatory inversion to explain phenotypic differences: Todesco *et al*. 2020) need to be complimented with an understanding of the range of SV types and their frequencies over a diverse range of different populations (e.g. Mérot *et al*. 2020), and studies of this range are scarce. Comparing different types of genetic variants will allow us to determine if SVs behave the same as SNPs within a population and will also shed light onto the underlying mechanisms of evolution.

In this manuscript, we seek to understand differences in patterns of genome-wide SNPs and SVs within the globally invasive European starling, *Sturnus vulgaris*. Starlings are native to the Palearctic and were introduced to Australia and North America in 1857 and 1890, respectively. Within Australia, starlings are distributed along much of the eastern and southern coastline, resulting from multiple geographically separated introduction sites and subsequent spread (Stuart *et al*. 2021b). We used whole genome sequencing of eight native range starlings from the United Kingdom (UK), eight invasive North American starlings (NA), and 33 invasive Australian (AU) starlings to examine patterns in genome-wide SNPs and SVs between populations and within Australia. Specifically, we aimed to assess overall SV patterns in starlings globally and compare the population genetic patterns of SNPs and SVs across the three starling populations. We then investigated signatures of SNP and SV adaptation between invasive AU and the native UK to assess whether variants diverging between the populations result from regional differences in balancing and directional selection, or have been maintained as neutral variants. We expect that balancing selection helps to maintain standing genetic variation (SNP and SV) within the native range (UK) onto which directional selection may act following introduction.

## 2. Materials and Methods

### 2.1. SNP Genetic sequencing data

The whole genome resequencing short read Illumina data used in this project are a combination of data from 24 individuals previously collected across the three continents of Australia (AU; invasive range), North America (NA; invasive range), and United Kingdom (UK; native range) (Hofmeister *et al*. 2021), and that of 25 additional individuals samples collected from sites across Australia (Supplementary Materials: Table S1). In total there were eleven sampling sites: New York, North America (eight individuals); Newcastle upon Tyne, United Kingdom (eight individuals); and Australia (33 individuals across nine sampling sites throughout Australia).

To obtain SNP variant information, the raw reads were processed using samtools (v1.9) (Li *et al*. 2009) and bedtools (v2.27.1) (Quinlan & Hall 2010), and adapters were removed using adapterremoval (v2.2.2) (Schubert *et al*. 2016). The processed reads were aligned to the starling reference genome *S. vulgaris* vAU1.0 (Stuart *et al*. 2021b) using bowtie2 (v 2.3.5.1) (Langmead & Salzberg 2012), and indexed using picard (v2.18.26) (*Picard toolkit* 2019). The gatk (v4.1.0.0) (Poplin *et al*. 2018) function *HaplotypeCaller* (GVCF mode) was used to call haplotypes for each sample, which were processed with *CombineGVCFs* and finally passed into *GenotypeGVCFs* to produce the initial VCF file (30,153,260 SNPs). The VCF file was put through an initial filter step in GATK (QD<2.0, FS>60.0, MQ<40.0, SOR>3.0), and a secondary filter using VCFtools (v0.1.16) (Danecek *et al*. 2011) (max proportion of missing alleles per SNP= 0.5, minor allele frequency = 0.03, min mean DP (approximate read depth) =2, max mean DP=50, min alleles=2, max alleles=2), producing a total of 19,256,335 SNP variants. These SNPs were then pruned in bcftools for linkage by removing sites with an r2 > 0.6 within 1000 bp site windows (Karlsson Linnér *et al*. 2019), resulting in 7,402,600 SNP variants. This created the variant file used in the local PCA and balancing selection analysis because these analysis require high density SNP data (Li & Ralph 2019; Siewert & Voight 2020). Hereafter, this dataset will be referred to as the whole genome SNP dataset (a brief summary of variant calling and data filtering for different analysis can be found at Supplementary materials: Fig. S1). For all other SNP analyses, high density was not required and having completely independent SNPs is more appropriate because it reduced overcontribution of groups of linked SNPs to admixture analysis, for example. We created a list of independent SNPs by thinning these variants in VCFtools, retaining only SNPs more than 5,000 bp from one another (--thin 5000), resulting in 186,205 SNPS. Hereafter, this dataset will be referred to as the thinned SNP dataset

### 2.2. SV calling using short-read data

We reanalysed the above 49 short-read WGS sample data for SVs using three different short read SV calling programs: lumpy-sv v0.2.13 (Layer *et al*. 2014), Delly v0.8.2 (Rausch *et al*. 2012), and Manta v1.6.0 (Chen *et al*. 2016). The three chosen SV callers use read-pair and split-read data (and in the case of lumpy-sv read-depth as well) as detection signals for the presence of SVs in sample read data. Further, calling SVs across multiple programs, and then combining the individuals calls into a consensus SV list based on overlap between datasets, helps to reduce the false positive rates from using a singular caller, which is of particular importance for short-read data which is more vulnerable to this type of error (Jeffares *et al*. 2017). For Lumpy-SV, we used the traditional pipeline; we mapped the read data using speedseq (Chiang *et al*. 2015), generated empirical size histograms and statistics for each sample BAM file using samtools (Li *et al*. 2009), identified SV’s by running lumpy on all samples simultaneously alongside the sample-specific histogram information, and used svtyper (Chiang *et al*. 2015) to call genotypes, with max reads set to 2000 (to remove regions of extreme coverage and decrease run time). For Delly, we followed the ‘germline SV calling’ pipeline, using the Bowtie2 mapped reads (see 2.1. SNP Genetic sequencing data). For Manta, we used the default pipeline using the Bowtie2 mapped reads (see 2.1. SNP Genetic sequencing data).

We then created a consensus SV file using Survivor v1.0.3 (Jeffares *et al*. 2017). We merged the three sets of SV calls for each sample using the parameters values of “1000 2 1 1 0 30” (max allowed distance between predicted break ends 1 kb, 2 SV caller support needed to retain a call, type and strand of the SV must agree, and minimum SV length of 30 bp). Precise definitions vary between studies, but in this study we define SVs as greater than 30 bp long. This created a consensus list of SV for each individual. We then merged across samples to obtain an overall SV file, using the parameters “1000 1 1 1 0 30” (same as above except minimum required support by caller was set to 1 to produce a merged SV list). This resulted in a file containing 20,083 SVs, and hereafter will be referred to at the unfiltered SV dataset. In all SV datasets, TRA (translocations) refers specifically to break ends.

### 2.3 SV repeat classification and final SV filtering

We characterised the repeat content of the unfiltered SV dataset by using RepeatMasker v4.0.7 (Smit *et al*. 2013) and a custom curated repeats database generated in Stuart *et al*. (2021a). We performed repeatmasker searches on all the of the SV sequences, using the first 30 bp of each SV sequence obtained by bedtools *getfasta*. Searching the repeat database for matches to the first 30 bp of each SV focused on the repeats that existed at the break end site, and ensured long SVs did not return numerous hits from the middle of their length.

We excluded any SV calls that were determined to be simple repeats (1,577 simple repeats, i.e. microsatellites as defined by RepeatMasker removed, see Fig. 1), because more erroneous SV artifacts are called in low complexity regions of the genome (Li 2014).

**Figure 1:**
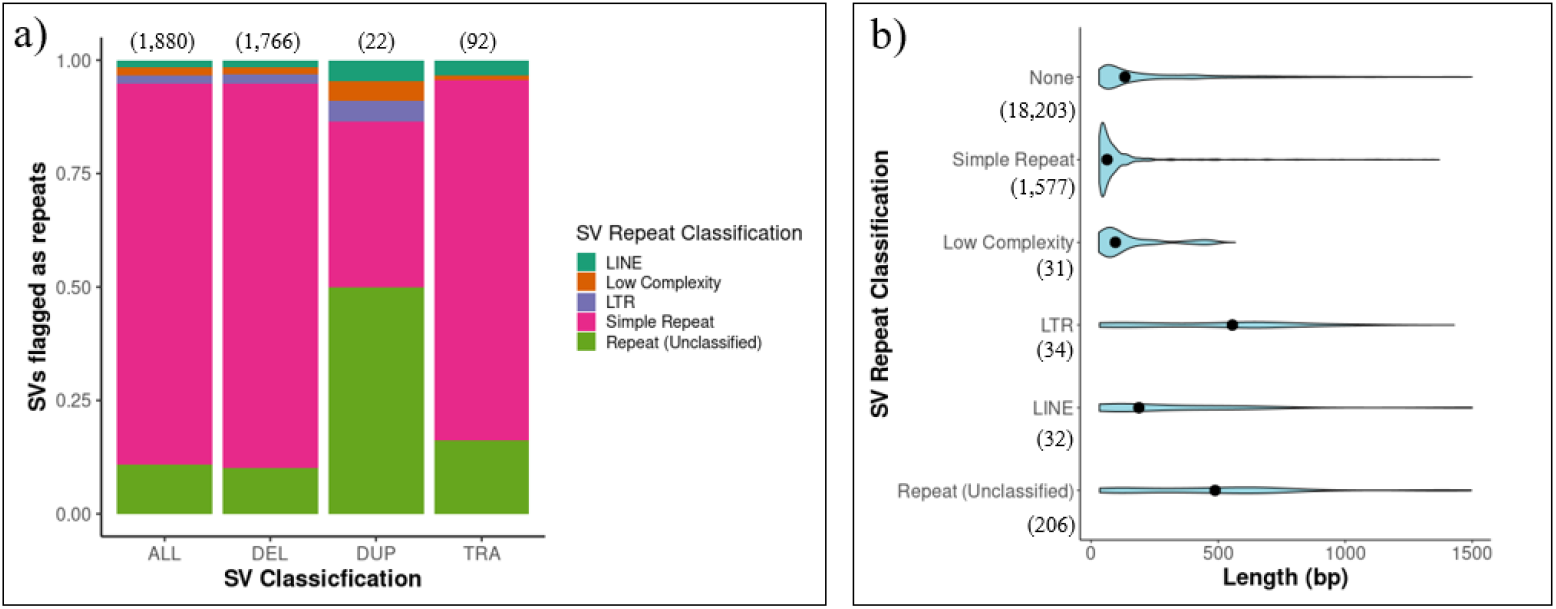
*Sturnus vulgaris* annotation of repeat elements in the unfiltered structural variant (SV) dataset across all 49 individuals. Panel (a) shows the proportion across all SVs, making up 9.36% (1,880/20,083) of the total length of the SV sequences. Panel (b) shows length distributions of the different SV repeat classes.

We then curated the SV file following recommended Survivor guidelines. We the assessed the Bowtie2 BAM files and computed the coverage of low mapping quality reads (MQ < 5) across each sample independently using samtools *view*. Survivor *bincov* was then used to produce a bed file of low-quality clustered regions per sample using the parameter values 10 5 (maximal distance between clustered regions, and only considering regions if coverage is larger than 5). These low-quality regions were then assessed across samples, and we removed SVs from the dataset if either or both ends of the SV overlapped with a low-quality clustered regions across more than 10 samples. This resulted in a file containing 15,543 SVs, and hereafter will be referred to at the quality-filtered SV dataset. The counts and size distribution of these SVs were then summarised, with vcftools used to calculate singletons.

The quality-filtered SV dataset was further filtered using VCFtools for missingness and MAF (minor allele frequency) at the same threshold as the SNP data (--max-missing 0.5 and --maf 0.03), resulting in 6,178 SVs. Hereafter, this dataset will be referred to as the popgen-filtered SV dataset, used for population structuring and diversity analyses, and directional and balancing selection analysis (Supplementary materials: Fig. S1).

To assess the impact of MQ and missingness thresholds, filtering was assessed by repeating the above MQ filtering across multiple thresholds (total individuals with low MQ coverage: 0, 5, 10, 15, 24, 37, 49), and the minor allele frequency densities plotted both with and without missingness filtering (--max-missing 0.5), and total SV counts summarised.

### 2.4 Examining SNP and SV density patterns

To investigate whether global patterns in SNP and SV density were correlated, Bedtools *coverage* was used to summarise SNP and SV counts (whole genome SNP dataset and quality-filtered SV dataset, Supplementary materials: Fig. S1) in 1 Mb bins along the 32 largest scaffolds of the starling genome, which was calculated for all samples, as well as within continental sample groupings (AU, NA, and UK). To assess whether SNP and SV density was correlated, the binned variant densities were analysed alongside each other in R v3.5.3 (R Core Team 2017) using the *glm*() function (family=quasipoisson), and residuals were retained. Bedtools *intersect* was used to find SVs that overlap with coding regions, against the starling genome annotation (Stuart *et al*. 2021b).

To check for any bias between total sequenced read counts in samples and their average genotype heterozygosity, the proportion of heterozygous variant sites for each individual was calculated for both SNPs and SVs (same files as above), and was assessed with a linear mixed model using the R function *lmer*() in the package LME4 v1.1-26 (Bates *et al*. 2015), with population set as a random factor.

### 2.5 Comparison of global SNP and SV population structuring and diversity

To contrast patterns of population structuring and genetic diversity, the following analysis was run on both the SNPs and SVs (thinned SNP dataset and popgen-filtered SV dataset, Supplementary materials: Fig. S1). Ancestry analysis was run using admixture (Alexander *et al*. 2009), while plink v1.9 (Purcell *et al*. 2007) was used to assess the major pattern of variance in the data via a principal component analysis (PCA). Nucleotide diversity (π) was assessed across the thirteen sample groupings using vcftools (eleven sample sites, plus one grouping of all AU sample sites, and an AU subset of the eight AU individuals from the same study as the NA and UK sample sizes (Hofmeister *et al*. 2021; Supplementary Materials: Table S1, individuals au01-au08). The program stacks v2 function *populations* (Catchen *et al*. 2013) was used to quantify the number of polymorphic markers, observed heterozygosity (Ho), expected heterozygosity (He), inbreeding coefficient (Fis) and private alleles for both SNP and SV genetic variants across these same sample groupings. Heterozygosity measures were assessed for statistical difference between sample sites using the R package ggstatsplot, using the *ggwithinstats()* function for nonparametric data (Patil 2021). Analysis of SV genetic diversity was conducted twice, once on the popgen-filtered dataset, and once on a dataset that omitted the MAF filtering to allow for rare variants (Supplementary materials: Fig. S1), to assess if measures of genetic diversity in SVs were biased against NA and UK because these sites had fewer sampled individuals than AU.

To investigate common unique SVs that had arisen in the AU population, the popgen-filtered SV dataset was filtered to retain only Australian-specific SV alleles not present in NA and UK (the 536 private allele SVs identified by stacks *populations*). We produced site frequency spectrum (SFS) plots for these Australian-specific SVs after dividing them into two groups; those that overlapped (either entirely or partially) with a gene, and those that did not. As a point of comparison, we also produced SFS plots for the popgen-filtered SV data set, and then subdivided this data set into three: SVs with no coding region overlap; SVs overlapping an entire coding region; SVs with one or more break ends overlapping a coding region.

The Australian-specific SVs were then further filtered to remove uncommon variants by filtering for a MAF of >0.15, which yielded 122 variants. We generated a list of candidate genes that exist within 1 kb of these common Australian-specific SVs using bedtools *intersect*, using the *S. vulgaris* vAU1.0 annotation (Stuart *et al*. 2021b).

To better understand the potential genetic impacts of the largest Australian-specific SV, we investigated the SV’s placement relative to SNP-based patterns of dissimilarity along the genome using local PCA, and plotted coding region maps of surrounding protein-coding genes. Local PCA uses high density SNP data (and so high genotyping certainty compared to SVs) to quantify genomic regions of similarity or dissimilarity to neighbouring regions along the length of a chromosome or scaffold using an ordination approach. By plotting MDS loadings along the length of a genome, it is possible to reveal regions of highly different population structure effects along the genome. Local PCA was conducted using the R package lostruct v0.0.4 (Li & Ralph 2019) on the chromosomes on which these variants were contained, in sliding windows of 1000 SNPs based on the whole genome SNP dataset (Supplementary materials: Fig. S1). Gene structure was mapped 10 kb upstream and downstream of the variants, and plotted using the R package sushi v1.31.0 (Phanstiel 2021).

### 2.6 Directional and balancing selection

We sought to investigate outliers that were possibly under directional selection in Australia versus the UK (i.e., variant sites that were divergent between the native UK and the invasive AU range). bayescan v2.1 (Foll & Gaggiotti 2008) uses allele frequency based approaches to identify genetic variants subject to natural selection by assigning a per-site posterior odds (PO, often plotted as log10 PO) probability estimated by comparison of explanatory models with and without selection. We conducted bayescan outlier analysis, with prior odds for the neutral model set to 10 for SNPs (-pr_odds 10), and 5 for SVs (-pr_odds 5, to reduce threshold for flagged outlier SNPs so that the strongest outlier variants would lie above the statistical threshold and be available for follow-up balancing selection analysis), and a false discovery rate (FDR) of 0.05. bayescan analysis was conducted using the thinned SNP dataset and popgen-filtered SV dataset, but with only UK and AU individuals (Supplementary materials: Fig. S1). Using bayescan we searched for signals of divergent selection between AU and UK, and to account for neutral high population substructure due to historic introductions demographic processes, the AU sample sites were split into two groups and the bayescan analysis conducted on each separately. The AU sample site subdivision was based on population substructure identified in Stuart & Cardilini *et al*. (2021) and was also corroborated by SNP based admixture patterns in this dataset (Fig. 3c below, AUeast group: Lemon Tree, Maitland, Dubbo, Hay; AUsouth group: Hay, Wonthaggi, Hobart, Meningie, Condingup, Munglinup; with Hay appearing twice due to ambiguous grouping, Supplementary materials: Table S1). PCA of all SNP and all SV outliers was generated using plink v1.9.

**Figure 2:**
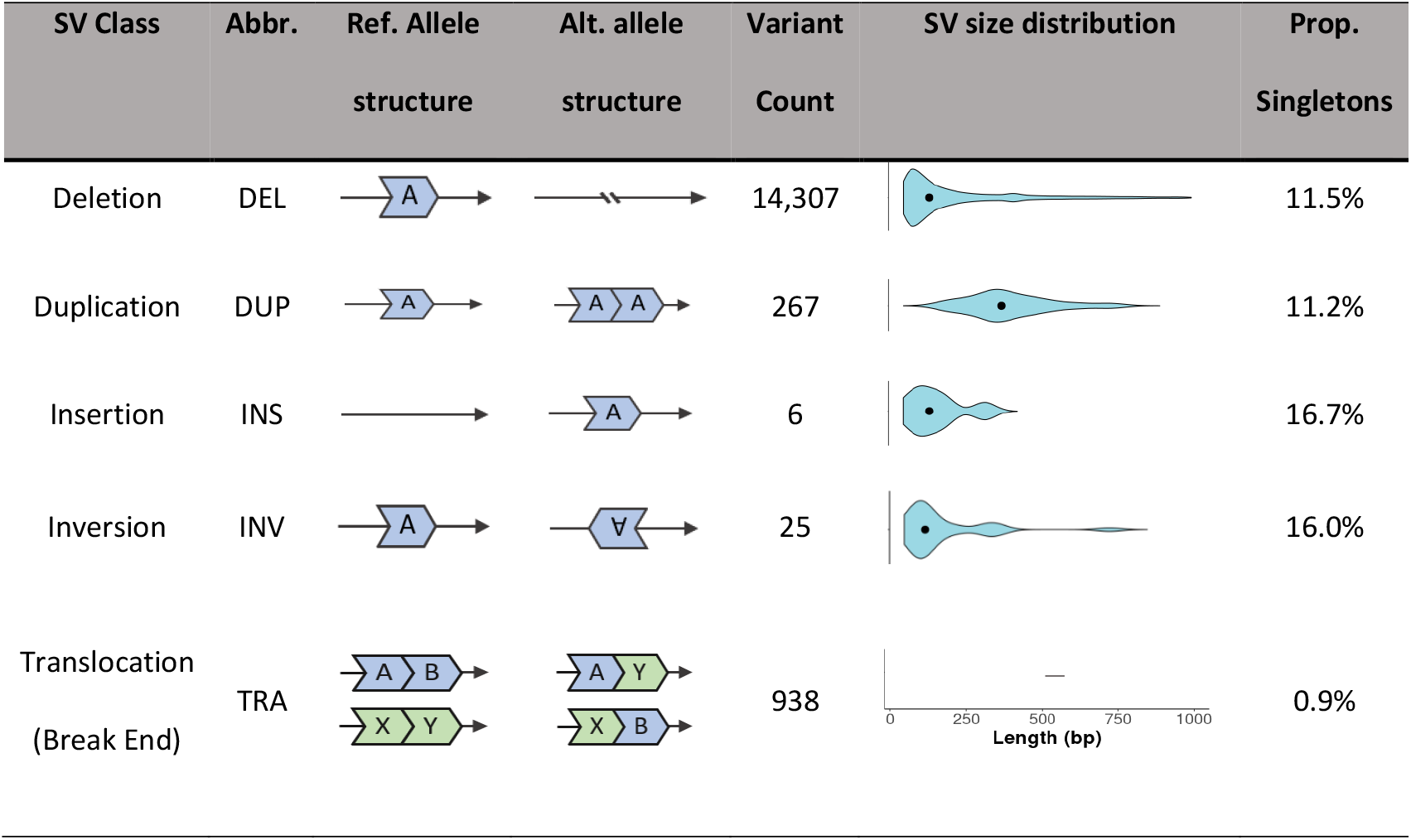
Summary of quality-filtered structural variants found across 49 *Sturnus vulgaris*. individuals spread across three continents, a total of 15,543 structural variants (SVs). SV size distribution black dot indicates median length of SV class. Proportion of singletons calculated using vcftools --singletons. SV size distribution not calculated for TRA as these SV were called break ends, and thus quantifying size across them would be inconsistent.

**Figure 3:**
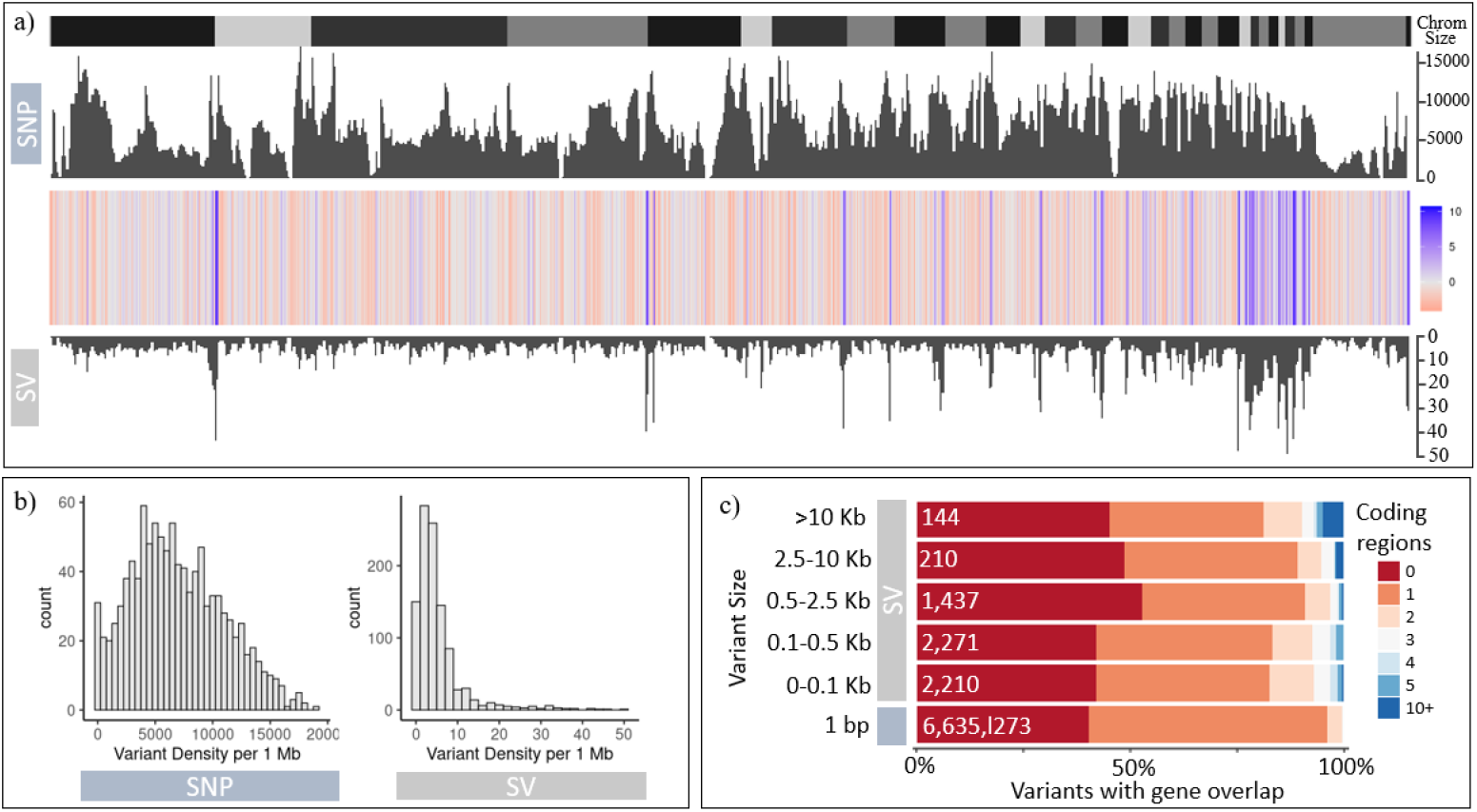
*Sturnus vulgaris* WGS single nucleotide polymorphism (SNP) and structural variant (SV) comparisons across the first 33 chromosomes. Panel (a) depicts the residuals from a regression of SV density on SNP density (10Mb windows) regression, with chromosome labels above the plot (positive residual blue, negative red). Panel (b) depicts a histogram of SV and SNP densities in 10 Mb windows. Panel (c) depicts proportion of each genetic variant size class that overlap with one or more genes, along with total variant count within each size classes displayed in white text. Some overlapped genes in the underlying annotation have resulted in 2 genes flagged for 1 bp (SNP) variant size class.

Balancing selection helps to maintain standing genetic variation within the native range and it has been hypothesised that this may provide the substrate upon which directional selection can act in invasive ranges (de Filippo *et al*. 2016; Huang *et al*. 2016; Koenig *et al*. 2019; Stern & Lee 2020). To test whether we found evidence for this in starling populations, we performed genomic scans for balancing selection on the UK SNPs and SVs (separately). Scans for balancing selection on SNPs were performed with the program betascan v2 (Siewert & Voight 2020), which uses relative allele frequencies of nearby sites to quantify a measure of balancing selection present at a loci, with high β^(1)^ scores being indicative of an excess of SNPs at similar frequencies within a region and thus balancing selection. Because this analysis relies on high density of SNPs, the whole genome SNP dataset was used, and filtered to only non-fixed alleles present in UK individuals (Supplementary materials: Fig. S1). Further, following recommended protocols (Siewert & Voight 2020), we only retained β^(1)^ scores for SNPs with a minimum MAF of 0.15 to reduce false positives (Siewert & Voight 2017) because balancing selection at low allele frequencies is unlikely to overcome drift and can exaggerate the effect of drift (Robertson 1962). Because β^(1)^ scores are generated using high density SNPs, SV data could not be used to directly generate β^(1)^ scores, nor direct β^(1)^ scores provided from SVs from SNP variants because SVs do not necessarily map directly to SNPs. Instead, we generated pseudo β^(1)^ scores for: INV by retaining all SNP β^(1)^ scores that overlapped with the SVs (+/-1000 bp); DEL, DUP, INS, TRA by retaining all SNP β^(1)^ scores that occurred within 1000 bp upstream of the first break end. The value of 1000 bp was chosen as this was the maximum allowed distance between SV calls during survivor consensus SV merging, and the different approaches for the SV types reflect our best estimations of how the underlying reference SNPs would best capture patterns in the overlaid SVs, and also because one may expect that patterns of allele frequencies may be similar for all SNPs immediately adjacent because these SNPs are expected to be linked. Then, for each SV, we obtained the trimmed mean (0.1 trimming, middle 90% data) for the β^(1)^ scores overlapping/occurring upstream of it, to try reduce the impact of outliers (we also tested a second approach, where β^(1)^ scores were not trimmed or averaged for each SV individually, but were simply pooled an used in the subsequent ANOVA). We used the r function *aov()* to test whether mean β^(1)^ scores differed across eight groups: SNPs under directional selection; SNPs not under directional selection; SVs (all DEL) under directional selection; and five SV types (DEL, DUP, INS, TRA, INV) not under directional selection.

## 3. Results

### 3.1. Patterns in *S. vulgaris* structural variants

Annotation of the unfiltered SV dataset identified repeat elements in 9.36% of the SV sequence break ends, with simple repeats making up the majority of the unfiltered repeat types (Fig. 1a). Repeat annotation profiles for DEL and TRA were very similar, while DUP variants contained a relatively smaller proportion of simple repeats than both these two groups (Fig. 1a). Simple repeats had the shortest length compared to other SV repeat classifications, while LTR and unclassified repeats (which contained a mix of species-specific repeats such as MITES) were the longest variant types (Fig. 1b).

Once simple repeats were filtered out, a total of 15,543 SVs (quality-filtered SV dataset) were identified over the 49 individuals in the short-read dataset, in comparison to the 7,402,600 unlinked SNPs identified in the whole genome SNP dataset. While the total number of SVs is much lower than that of SNPs, the total SV length (26,443,169 bp) covers 3.57x the total length of the identified SNP variants.

Across these SVs, DEL were the most common SV class, followed by TRA and DUP (Fig. 2). A very small number of INV and INS were detected. Of the different SV classes, DUP were on average the largest (median length = 359) with the other SV size classes having roughly similar sizes (Fig. 2; median lengths 117, 117, and 101 for DEL, INS, and INV respectively; no sizes provided for TRA because break ends have undefined sizes). The proportion of singleton variants across all SV classes was low (16.7% or less), particularly from TRA (0.9%) (Fig. 2).

The density of the SNPs and SVs were highly correlated (general linear model: t value = 13.54, p value < 0.0001). Visualisation of residual patterns between SNPs and SVs revealed that for most of the length of the genome, SVs are underrepresented; however, there is SV enrichment near the end of chromosomes (Fig. 3a; enrichment is also visible on smaller chromosomes, though this is likely a result of having a higher proportion of the chromosome length as chromosome ends). Overall, SVs occurred in much lower density (most commonly 6 per 10 Mb) than SNPs (most commonly 5,000 per 10 Mb) (Fig. 3b). SNP and SV density patterns were very similar across all three populations (Supplementary materials: Fig. S2). Analysis of patterns of coding region coverage for variants of different sizes indicate that roughly 80-90% of variants overlapped one or no coding regions, though as SV size class increased so too did the number of SVs containing 5 or more coding regions (Fig. 3c).

Regardless of the MQ threshold used to filter SVs, the minor allele frequency kernel density estimates were very similar (Supplementary materials: Fig. S3). Missingness filtering however did have a major impact on the minor allele frequencies, severely reducing the number of SVs at intermediate allele frequencies. Finally, no significant relationship was found between individual heterozygosity proportions and total sequence reads (Supplementary materials: Fig. S4).

### 3.2. Population genetics of *S. vulgaris* using whole genome SNP and SV data

Population structure analysis conducted using admixture on SNPs and SVs supported similar ancestry patterns across sampled populations, though SV based population structuring contained more signals of mixed ancestry within Australia particularly at higher K (number of ancestral subpopulations) values (Fig. 4c, Supplementary materials: Fig. S8). Both datasets supported similar ancestry patterns for NA and UK. Meanwhile, AU populations contained mixed ancestry signals, with the westernmost range-edge site (Munglinup) remaining distinct. A very small amount of AU subpopulation structuring was identified across AU sample sites, though this pattern was not apparent in the SV dataset. Because the K values 1-3 received similar cross-validation error support (Supplementary materials: Fig. S9), we plotted K=3 as the highest K value allowing us to discern more patterns, and also as means of comparing to similar analysss completed previously using some of these samples (Hofmeister *et al*. 2021: APPENDIX F).

**Figure 4:**
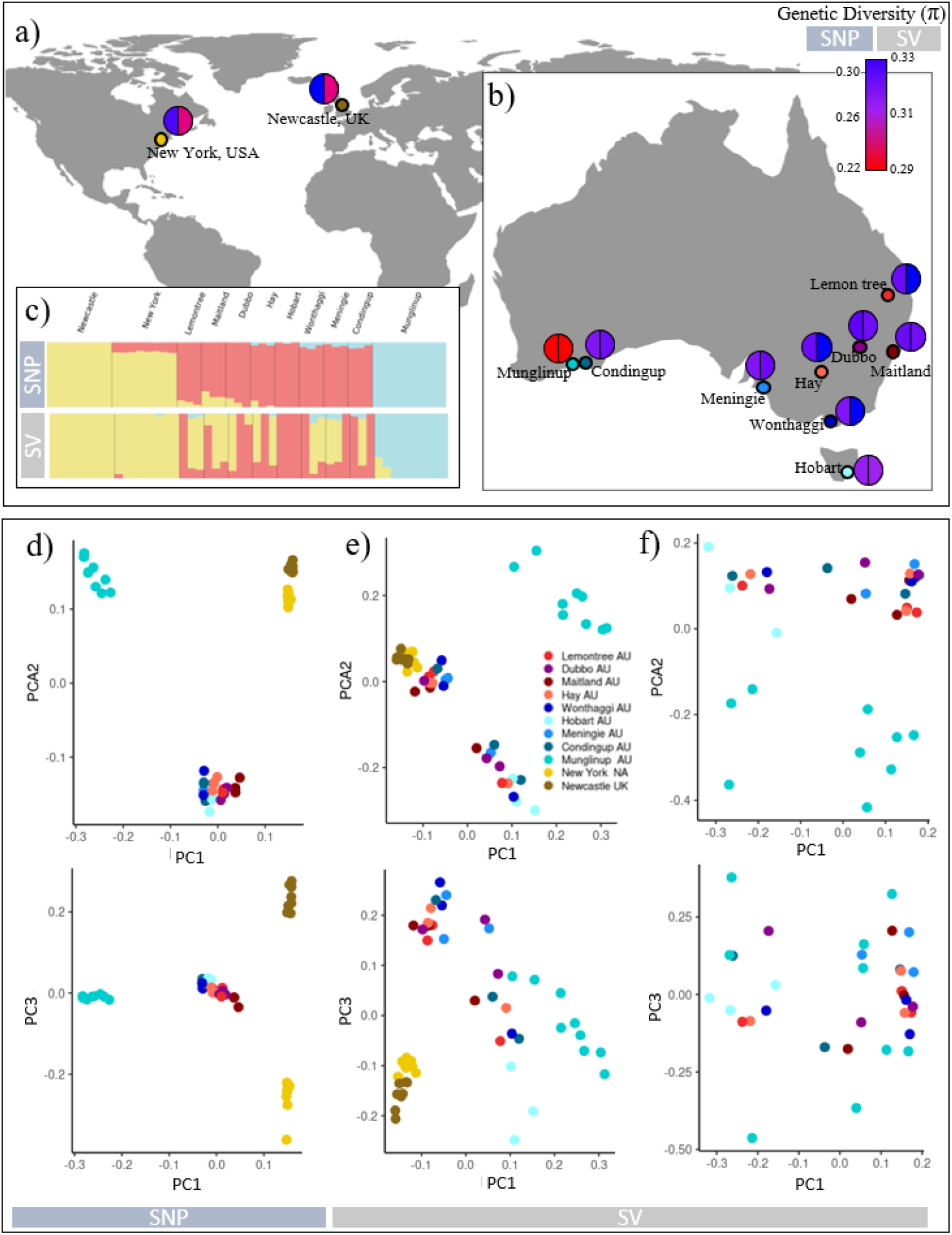
*Sturnus vulgaris* WGS single nucleotide polymorphism (SNP) and structural variant (SV) population structure and genetic diversity. Panel (a) and panel (b) indicate sampling sites, and colours represent genetic diversity (π) at each locality (left is SNP, right is SV; π variance and standard error information available in Table S2). Panel (c) contains admixture plots from SNPs (top) and SV (bottom). Panel (d) contains PCAs of SNP variants (thinned SNP dataset, n=186,205), panel (e) contains PCAs of all SVs (popgen-filtered SV dataset, n= 6,178), and panel (f) contains PCA of all common Australian-specific SV allele variants (n=122).

Similar to the admixture analysis results, both SNP and SV PCA analyses supported the clustering of NA and UK individuals, as well as distinctly separate clustering of the western range-edge from the rest of the Australian sampling sites (Fig. 4d&e). Finally, PCA analysis revealed some artifactual clustering in the SV data for AU samples sequenced at a lower depth (Fig. 4 e; individuals clustering closest to NA and UK had lower sequencing depths, Supplementary Material: Table S1). While overall heterozygosity was not correlated with sequencing depth (Supplementary materials: Fig. S3), a subset containing the eight AU individuals with similar sequencing depths (to UK and NA) was included in the genetic diversity analysis to ensure data was not biased against UK and NA individuals (which had smaller sample sizes than Australia), and was largely in congruence with other AU genetic diversity results (Supplementary Material: Table S3).

Comparing genetic diversity patterns across the global sampling sites, we found that patterns of diversity in SVs were broadly similar to SNP diversity patterns, with some key differences. We found that overall SNP genetic diversity (π, Ho, He, Fis) was higher in the NA and UK populations compared to AU (Fig. 4a & b, Table 1, Supplementary Materials: Table S2, Table S3). However, when considering SV genetic variants, the genetic diversity across these populations was more similar, and in the case of He was often higher in AU compared to NA and UK. While these differences are slight, many pairwise comparisons were significantly different (Supplementary Materials: Fig. S5-7). This shift in the relative genetic diversities in UK, NA, and AU was still apparent even in the absence of MAF filtering; because MAF filtering filters out rare variants, this may bias estimates in the two populations with smaller sample size (Table 1). Both SNP and SV genetic diversity patterns revealed that Munglinup contained minimal genetic diversity relative to the other AU sample sites (Fig. 3b). Despite this reduced genetic diversity, Munglinup was found to have many private alleles (105 for the thinned SNP dataset and 12 for the popgen-filtered SV dataset) in comparison to the Hofmeister et al. subset of a similar number of AU individuals (56 for the thinned SNP dataset and 4 for the popgen-filtered SV dataset).

**Table 1:** Single nucleotide polymorphism (SNP) and strucutral variant (SV) summary statistics for *Sturnus vulgaris* across study sample sites and continents. Ho = Observed Heterozygosity, He = the within population gene diversity, Fis = inbreeding coefficient, No. polymorphic site indicates the number of nucleotide positions that are polymorphic in at least one indivdual within a sample site or sample grouping. Number of polymorphic sites, Ho, He, and Fis were assessed using stacks *populations*. Variance and standard error information available in Table S3. SNP dataset was the thinned SNP dataset (186,205 SNPs), and SV datasets were the popgen-filtered SV dataset (6,178 SVs), and an alternate version of this that had no minor allele frequency (MAF) filtering (7,898 SVs).

| Continent | Sampling Site (sample number) | SNP (186,205) |  |  |  | SV (6,178) |  |  |  | SV (7,898) |  |  |  |
| --- | --- | --- | --- | --- | --- | --- | --- | --- | --- | --- | --- | --- | --- |
|  |  | No. Polymorphic Sites | Ho | He | Fis | No. Polymorphic Sites | Ho | He | Fis | No. Polymorphic Sites | Ho | He | Fis |
| **Australia**<br>*Invasive* | All sites (33) | 167,868 | 0.279 | 0.234 | 0.029 | 6,105 | 0.327 | 0.312 | -0.004 | 6,623 | 0.259 | 0.247 | -0.002 |
|  | Lemon Tree (3) | 111,778 | 0.293 | 0.242 | -0.006 | 4,126 | 0.340 | 0.266 | -0.014 | 4,198 | 0.269 | 0.211 | -0.011 |
|  | Maitland (3) | 111,813 | 0.291 | 0.242 | -0.002 | 3,940 | 0.321 | 0.255 | -0.004 | 3,998 | 0.254 | 0.201 | -0.003 |
|  | Dubbo (3) | 111,928 | 0.285 | 0.242 | 0.009 | 4,081 | 0.333 | 0.263 | -0.009 | 4,179 | 0.264 | 0.209 | -0.007 |
|  | Hay (3) | 112,122 | 0.295 | 0.242 | -0.007 | 4,070 | 0.332 | 0.263 | -0.007 | 4,139 | 0.262 | 0.208 | -0.005 |
|  | Wonthaggi (3) | 107,553 | 0.280 | 0.234 | 0.002 | 4,165 | 0.338 | 0.269 | -0.006 | 4,254 | 0.268 | 0.214 | -0.004 |
|  | Hobart (3) | 105,256 | 0.274 | 0.228 | -0.001 | 3,849 | 0.320 | 0.249 | -0.013 | 3,913 | 0.253 | 0.197 | -0.011 |
|  | Meningie (3) | 107,729 | 0.277 | 0.233 | 0.005 | 4,007 | 0.325 | 0.258 | -0.005 | 4,063 | 0.256 | 0.204 | -0.003 |
|  | Condingup (3) | 107,024 | 0.284 | 0.232 | -0.009 | 3,976 | 0.328 | 0.259 | -0.007 | 4,034 | 0.259 | 0.204 | -0.006 |
|  | Munglinup (9) | 111,933 | 0.236 | 0.212 | -0.027 | 4,795 | 0.322 | 0.271 | -0.054 | 4,886 | 0.253 | 0.213 | -0.042 |
|  | Hofmeister et al. Subset (8) | 146,766 | 0.286 | 0.272 | 0.011 | 5,278 | 0.333 | 0.291 | -0.027 | 5,425 | 0.263 | 0.230 | -0.021 |
| **North America**<br>*Invasive* | New York (8) | 154,991 | 0.311 | 0.289 | -0.006 | 5,023 | 0.328 | 0.277 | -0.047 | 5,228 | 0.260 | 0.220 | -0.036 |
| **United Kingdom**<br>*Native* | Newcastle upon Tyne (8) | 158,574 | 0.315 | 0.295 | -0.002 | 5,082 | 0.330 | 0.279 | -0.048 | 5,310 | 0.262 | 0.222 | -0.037 |

Comparing patterns of genetic diversity between SNPs and SVs revealed that overall, both Ho and He estimates were higher using SV data than from the SNP dataset. The highest levels of SNP He and Ho were found in Australia in Lemon Tree and Hay, while the highest levels of SV He and Ho were found in Lemon Tree and Wonthaggi. Across all sampling sites, we found that Ho was higher than He for both the SNP and SV dataset (though the difference was more extreme in the SV dataset). This was also reflected in Fis estimates which were high from SNP data of AU samples but not in the SV data from these individuals (Table 1).

### 3.3. Australian-specific *S. vulgaris* SV’s

A total of 536 Australian-specific SV were found across the 33 Australian samples. The SFS of the Australian-specific SVs that overlapped (either partially or completely) a coding region and those that did not were very similar (Fig. 5a&b), with a vast majority of the variants falling below MAF levels of 0.1. The SFS profile of these SVs were quite different from the popgen-filtered SVs, with a stronger bias towards more common (higher MAF) alleles (Fig. 5c). For popgen-filtered SVs with break ends overlapping genes, we observed a comparatively more even frequency distribution, with more variants found to have intermediate frequencies (Fig. 5e) compared to popgen-filtered SVs that did not overlap genes (Fig. 5d), and for popgen-filtered SV with complete coding region overlap high MAF levels were most common (Fig. 5f) (though this grouping of variant was considerably more rare overall).

**Figure 5:**
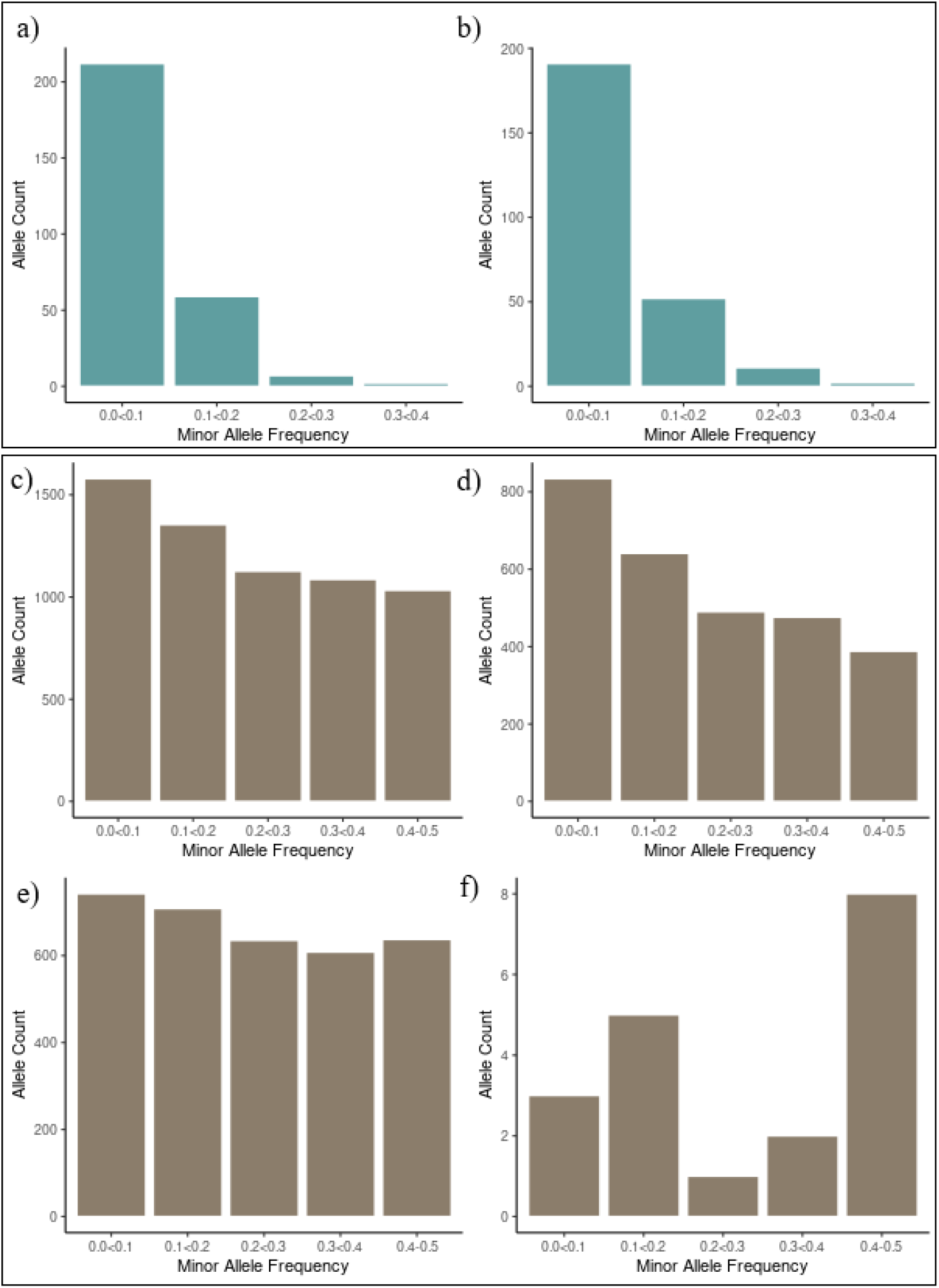
Folded site frequency spectrum (SFS) for structural variant (SV). Panel (a) and (b) depicts SFS of Australian-specific SVs with no coding region overlap and with (either partial or complete) coding region overlap respectively. Panel (c) depicts the SFS of the full popgen-filtered SV data set, panel (d) the SFS of popgen-filtered SV with no coding region overlap, panel (e) the SFS of popgen-filtered SV with break end coding region overlap, and panel (f) the SFS of popgen-filtered SV with complete coding region overlap.

When these 536 Australian specific SVs were filtered for common variants (a minimum MAF og 0.15), a total of 122 variants were retained. We observed that the strong distinction between range-edge AU and the rest of the sample sites was minimised but still somewhat present when PCA analysis was conducted only on the 122 common Australian-specific SV alleles (Fig. 4f). These variants were mainly deletions (as found in the overall SV dataset), and ranged from 30-25,098 bp in length. Of the variants greater than 1 kb in length, roughly half were found to overlap or be within 1 kb of coding regions (Supplementary materials: Table S4). The largest three of these were plotted with respect to their SNP based local PCA plots (Supplementary materials: Fig. S11). While none of these variants aligned with major signatures of genomc architechture seen in the local PCA plots (e.g. MDS peaks), all three were found to exist at points in the genome where patterns of SNPs within the SV were relatively dissimilar from surrounding SNP patterns, lending support to these being legitimate SVs (i.e. all three red lines passed through points in the plots where the dots, representing windows of 1000 SNPs, were found to be quite different respective to their nearest neighbouring points in terms of their loading on either MDS coordinate 1 or 2; Supplementary materials: Fig. S11).

A wide variety of coding regions underlying a wide range of biological processes were identified within the Australian-specific SVs (Table S4, Fig. S10). For the largest three variants, a 25 kb DEL, overlapped the gene *Low-density lipoprotein receptor-related protein 1B* (*LRP1B*) and one protein of unknown origin, and occurred downstream of *Nuclear receptor subfamily 4 group A member 2* (*NR4A2*) (Supplementary materials: Fig. S11a). The second largest variant, an 18 kb deletion, occurred upstream of *CD7 molecule* (*CD7*) (Supplementary materials: Fig. S11b). The third largest SV was a 9 kb duplication over two *Argininosuccinate lyase* genes (*ASL* and *ASL2*), and was upstream of Transmembrane Protein 132E (*TMEM132E*) (Supplementary materials: Fig. S11c).

### 3.4. Signatures of SV adaptation globally and within Australia

Out of the 186,205 SNPs tested for signatures of directional selection between AU and UK using bayescan, we identified a total of 52 SNPs (East: 3; Fig. 6a, South: 49; Fig. 6b). Out of the 6,178 SVs tested for signatures of directional selection using bayescan, we identified a total of nine SVs (East: 2; Fig. 6d, South: 7; Fig. 6e). PCA analysis of these outliers revealed much greater diversity in UK compared to AU for the outlier SNPs flagged as under putative directional selection between AU and UK (Fig. 6c), however the opposite was true for outlier SVs, which showed much greater diversity across AU samples (Fig. 6f) (although the sample size for this last group was small and therefore may have been affected by data artifacts). Similar to the PCA plot of common Australian-specific SV alleles (Fig. 4f), differentiation between the range-edge (Munglinup) samples and the rest of AU was heavily reduced in both SNP and SV outlier PCA plots analyses compared to analyses of neutral SNPs (Fig. 4 d&e).

**Figure 6:**
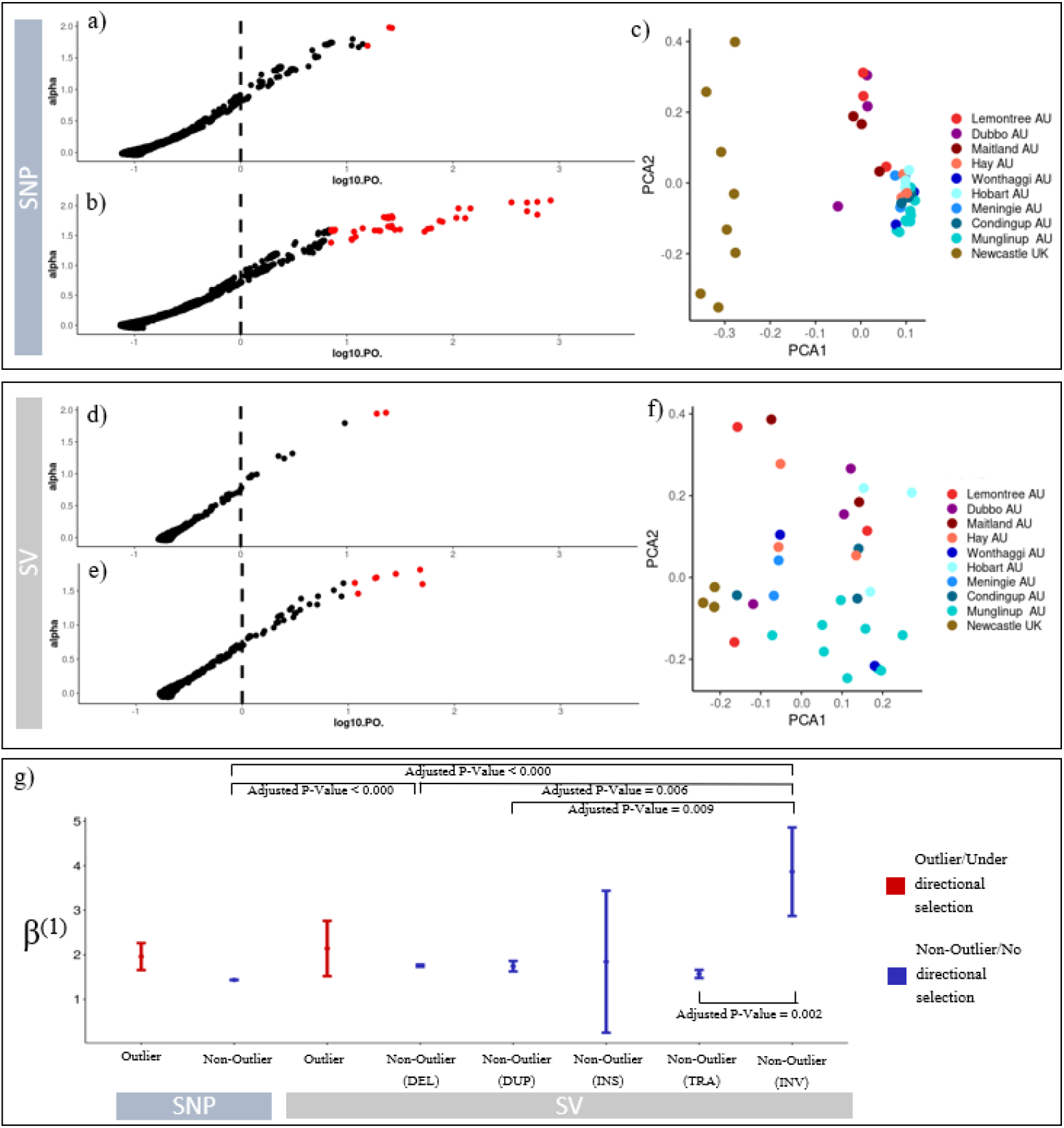
Genetic variants under putative directional selection in *Sturnus vulgaris* across the native range and invasive Australian range and assessment of balancing selection, for both single nucleotide polymorphisms (SNPs) and structural variants (SVs). For SNP variants, panel (a) depicts bayescan results for the UK-AUeast test for loci under balancing selection, with alpha (locus-specific component of selection) plotted against log_10_PO (log of posterior odds) values and flagged outliers indicated in red. Panel (b) similarly depicts bayescan results, but for UK-AUsouth comparison. Panel (c) depicts a PCA of all outlier SNPs (pooled across the UK-AUeast and UK-AUsouth analysis) for all AU and UK individuals. Panels (d), (e), and (f) depict the same as above, but for SVs. Panel (g) depicts mean β^(1)^ scores (balancing selection within the native UK range) for SNPs and SVs under directional selection (bayescan outliers between AU and UK) and no directional selection (+/- standard error) in UK individuals, alongside the adjusted p-value of posthoc Tukey’s pairwise tests of significance. SV pseudo β^(1)^ scores calculated by obtaining a trimmed mean (0.1 trimming) of overlapping/upstream SNP β^(1)^ scores for each SV individually, before averaging this score within each SV type group.

Comparing outliers (under directional selection) to non-outlier loci across SNPs and the different SV types (DEL, DUP, INS, TRA, INV), we found a significant difference between average β^(1)^ scores (levels of balancing selection within the native range) in sample groupings (F_7, 113659_ = 26.76, p-value < 0.0001; Fig. 6g; Supplementary Materials: Table S5). Post hoc analysis using Tukey’s pairwise test revealed that non-outlier SNPs had statistically lower β^(1)^ scores (lower levels of balancing selection in the native range) than non-outlier SV DELs and INVs, while non-outlier SV INVs had statistically higher β^(1)^ scores than non-outlier SV DELs, DUPs, DUPs, and TRA (Supplementary Materials: Table S5). Using the second approach to obtaining pseudo β^(1)^ scores yielded similar patterns across tested SNP and SV groups, but with more significantly different pairwise comparisons (due to larger data counts) (Supplementary Materials: Fig. S12, Table S6).

## 4. Discussion

SVs are an integral component of the genetic diversity within a species, with the potential to have profound functional consequences for an organism (Puig *et al*. 2004; Todesco *et al*. 2020; Hämälä *et al*. 2021). Clarifying the roles that SVs play in allowing an invasive species to persist and evolve within their newly invaded range is an integral next step to understanding how genomic architecture enables invasive populations to flourish. The findings of our research demonstrate that even within recently diverging lineages or populations, there may be a great deal of structural variation, either retained from the ancestral pool of variants, or recently derived within the newly isolated population. Further, patterns of SV genetic diversity do not necessarily reflect patterns in SNP diversity, either when considering interpopulation diversity patterns, or patterns of diversity across the genome. Finally, there is apparent directional selection within the invasive Australian range that appears to be enabled by the standing genetic variation introduced in founding individuals, and may have been maintained within the native range by balancing selection.

### 4.1 Contrasting SNP and SV patterns within the genome

Broadly, patterns in SNPs and SVs were found to be relatively similar across the three populations of starlings investigated in this study. Regarding the overall number of SNPs and SVs, we found that SV density characterised with our short-read data was similar to that of avian corvid species (Weissensteiner *et al*. 2020), though surprisingly the whole genome SNP density was quite high in comparison to other species, although this maybe also due to different sampling schemes or filtering parameters choices.

We found that densities of *S. vulgaris* SNPs and SVs were significantly correlated, though identified an increase in the densities of SVs near the ends of chromosomes. This may in part be due to artifacts, for example assembling repeat regions in reference genomes is difficult (Treangen & Salzberg 2011) and thus may result in underlying reference mistakes onto which the reads will try to map. However, research on human SVs has identified that centromeres and, more commonly, subtelomeric regions and telomeres are genomic regions where high proportions of SVs are observed (Collins *et al*. 2020), which (even accounting for repeat filtering step) provides a biological explanation for these results.

We found that even in structural variants up to lengths of 10 kb, the proportion of variants overlapping coding regions was similar to that of SNPs (each a single base pair in length). Characterisation of standing structural variants in inbred lines of the plant species *Mimulus guttatus* (Flagel *et al*. 2014) found that SV indels overlapping with coding region regions were rarer than that would be expected by chance, with this high prevalence of non-coding and specifically repeat regions being seen in the profile of other avian species’ standing SVs (Weissensteiner *et al*. 2020). While we did filter out approximately 9% of the SVs that were identified as simple repeats, SV datasets are often repeat heavy and likely our repeat analysis only annotated a proportion of the true repeat (given comparisons to other SV analysis which report repeat overlap of up to 80% for called SVs ; He *et al*. 2019; Penso-Dolfin *et al*. 2020; Weissensteiner *et al*. 2020).

It is perhaps because of this bias of SVs residing in non-coding regions that we found an unexpectedly high proportion of SVs at intermediate frequencies. This high proportion of SV heterozygotes may be in part a result of some short read structural variant callers, particularly Delly, incorrectly genotyping homozygous non-reference genotypes as heterozygous (Kronenberg *et al*. 2015). It may be interesting to note here that because of this bias against calling homozygous non-reference genotypes correctly, we might expect to find an increase in SV observed heterozygotes (Ho) in UK and NA birds (as these would be more genetically distinct from the AU reference genome), however, this was not the case (Table 1) and thus likely did not impact our results. Currently, one of the greatest challenges faced when calling SVs is the difficulty and low confidence of calling an individual’s genotype for an identified SV (Chander *et al*. 2019). This high proportion of SVs at MAF levels of 0.5 was heavily reduced during missingness filtering which presumably selects against SVs that are hard to call and genotype (Supplementary Materials: Fig. S3), but nevertheless there was still a surplus of heterozygotes than would be predicted based on allele frequencies across SV datasets when compared to SNPs (Table 1). While increased heterozygosity may be artifactual, this overall enrichment for heterozygosity may also have biological underpinning, because the bias for SV occurrence in non-coding regions means a reduction in processes which reduce mutation accumulation rates in protein coding regions, such as purifying selection (Gorlov *et al*. 2006) or mismatch-repair (Frigola *et al*. 2017). An alternative viewpoint suggests that we may expect SV allele frequencies to be lower due to the potentially deleterious effects that may have on phenotype, owing to their increased size and hence increased likelihood of functionally altering the genome (Kou *et al*. 2020). Studies have found that relative SV allele frequencies may be either low (Kou *et al*. 2020) or high (Flagel *et al*. 2014), and undoubtedly the standing SV load that can be maintained in a population is complex, and dependent on taxa or even population specific selection regimes.

### 4.2 Contrasting SNP and SV patterns across populations of a recent invasive species

Genetic structure assessed using PCA and admixture analyses found similar results across the SNP and SV data for AU, UK and NA individuals. AU individuals are genetically quite distinct from NA and UK (which both clustered together) across both SNP and SV both datasets. This clustering pattern was previously identified in some of the individuals used as part of this study (Hofmeister *et al*. 2021: APPENDIX F), but importantly our dataset expanded the AU sampling scheme and therefore provides important perspective on the extreme levels of genetic divergence at the westernmost AU range-edge (Munglinup) using both SNPs or SVs. Previous studies have demonstrated that SV genotyping may have many false positives and negatives (Chakraborty *et al*. 2019; Weissensteiner *et al*. 2020). Technical difficulties such as the variant may span the sequencing molecule’s length many time over, the repeat content within the variant, or the overall complexity of the rearrangement, means that short-read sequencing in particular is prone to SV genotyping errors (Chander *et al*. 2019; Ho *et al*. 2020; Wold *et al*. 2021). Hence it is reassuring to find that broad patterns of genetic variation were quite similar across the high confidence SNP and lower confidence SV dataset (Fig. 4).

Previous studies using microsatellite, mitochondrial sequence, and reduced representation sequence data have identified high genetic differentiation at the western-most range-edge of Australian starlings (Rollins *et al*. 2009, 2011; Stuart *et al*. 2021a). This result is reflected too in our whole genome resequencing SNP and SV data. Considering the close proximity of Munglinup to the nearest other sampling site (Condingup), the genetic diversity loss as well as occurrence of unique genetic variants in both SNPs and SVs in Munglinup is interesting. Patterns of genetic variants identified at this range-edge highlights the need for further native range sampling to establish whether the genetic variants private to this site are novel mutations or whether an undetected introduction from a different (unsampled) part of the native range has occurred in Western Australia, because so far the novel mitochondrial haplotype identified in this region remains unique (Rollins *et al*. 2011; Bodt *et al*. 2020). Interestingly, this range-edge differentiation was lost when analysis was restricted to just SNPs and SVs variants under directional selection between AU and UK (Fig. 6), and was somewhat reduced though still evident in the common Australian-specific SV allele (Fig. 4f). This may be indicative of different processes driving adaptive and neutral genetic SV patterns, with the adaptive processes being driven by similar selection regimes across the whole introduced range, reducing the range-edge effects.

Heterozygosity measures across SNPs and SVs revealed a difference in genetic diversity patterns (Ho, He, Fis) across some sampling sites in our data, despite the above discussed statistical correlation between SNP and SV densities. SNP data indicate a reduction in genetic diversity following introduction to Australia, whilst genetic diversity estimated from SVs is highest in Australian starlings. The relatively higher diversity of SVs (compared to patterns of SNP diversities between continents) in the AU range is not due to MAF filtering, because this pattern remains even when rare alleles were included. Importantly, our SV quality filtering processes filtered out SVs that overlapped with regions of low mapping quality (calculated on an individual basis). Where a read fell below the quality threshold, it was removed across all individuals, which should have minimised biases against the smaller sized NA and UK sample groups. Follow up analyses with long read data or improved bioinformatic SV detection methods may help validate whether these results hold true for all types of SVs, or may reveal that these results only apply to the classes of SVs that are more easily detected by short-read SV callers, like deletions and duplications. Further, while simple repeats were filtered out for this study as they are less likely to be genotyped correctly (Li 2014), long read sequencing will allow these and other repeat-based genetic variants (for example transposable elements) to be better characterised and explored. Finally, generation of more high quality genomic resources for this species, including a global pangenome, will enable the exploration of the role reference bias may play in influencing these results (Chen *et al*. 2021).

It seems apparent that at least for the SVs detected using our approach, AU individuals contain greater diversity than NA and UK individuals. Higher SV diversity in Australian starlings could be a result of admixture of the multiple introductions into Australia. However, under these circumstances, we would expect SNPs to also then reflect the higher diversity. Possibly, SV diversity is maintained through some demographic or evolutionary means; it has been found in humans that there is an overrepresentation of SVs in genes that are important for interacting with the surrounding environment (e.g. olfaction and response to external stimuli; Feuk *et al*. 2006a, b; Nguyen *et al*. 2006). Difference in SV and SNP allele frequency patterns have been found across multiple Asian rice varieties (Kou *et al*. 2020), thought to result at least in part from selection via domestication. Examination into how and why some populations have differing levels of genetic diversity across SNPs and SVs (and other forms of genetic variation) is integral for a better understanding of how genetic diversity is maintained within a population. As structural variant calling approaches improve and become more standardised across studies, investigation of broadscale patterns of relative SNP and SV densities across taxa will be useful. These studies can address questions about how mutational load may contribute to future population adaptation, despite the potential to confer deleterious effects on individuals (Wagner *et al*. 2017).

As would be expected of newly arisen variants, most of the Australian-specific SV alleles were rare (Fig. 5a&b), which contrasted strongly to the less variable SFS of the popgen-filtered SV data set, and subsets. Within SVs with at least one break end overlapping a gene, we may expect a SFS preferencing lower MAF based on the potentially deleterious nature of the break ends impact on coding region function, however contrary to this we see higher MAFs are still relatively common (Fig. 5e). It is possible that some of these SVs were small enough to be contained entirely within the same coding region and therefore have a non-negative (or even positive) effect coding region function and hence selection, a supposition supported in part by the SFS of SVs containing entire genes showing even more frequent high MAFs (Fig. 5f). Overall, the SFS plots indicate that global starling SVs with potentially functional effects are being maintained within the population; contrasting sharply to the much newer and rare Australian-specific SVs.

While most of the common Australian-specific alleles were smaller than a few hundred base pairs in length, substantially longer ones were observed, and many overlapped with coding regions that may feasibly play a role in AU specific starling evolution. For example, there may be a potential role for selection at *TAS2R40* in response to taste perception of novel food sources (Behrens *et al*. 2014), and IKZF1 in lymphoid differentiation and hence immune response (Marke *et al*. 2018). In particular, the third largest common Australian variant (9 kb duplication on chromosome 19) was found to overlap the two paralog genes *ASL* and *ASL2*, themselves orthologs of the gene ASL that plays a direct role in the urea cycle and ammonia detoxification in humans (Wertman 2012). While birds do not have a urea cycle as they who convert ammonia straight to uric acid, these genes still play an important role in the generation of the amino acid arginine and in lens formation (Piatigorsky & Wistow 1989; Erez *et al*. 2011). That being said, we observed much variety in MDS 1 and 2 loadings along the focal chromosomes, with strong peaks and troughs possibly indicating large scale INV or TRA, an indication that there is much more to be explored via long-read sequencing regarding the overall genomic architecture of this species.

### 4.3 patterns of selection in *Sturnus vulgaris* SNPs and SVs

While environmental conditions may elicit directional selection on genetic variants, under fluctuating or intermediate environmental conditions balancing selection may help to maintain genetic variation within populations, countering the tendency for polymorphisms to be lost via genetic drift. We found that that the levels of native range balancing selection were significantly higher in some groups of non-outlier SVs than non-outlier SNPs, suggesting balancing selection may play a greater role in maintain moderate SV allele frequencies compared to SNPs. We also note that non-outlier INVs had the highest levels of balancing selection, which may indicate that these types of SVs in particular are prone to persist under balancing selection. While the aforementioned technical difficulties with SV genotyping may influence the identification of SVs during divergent selection analysis, the β^(1)^ scores for SVs were calculated directly from underlying SNP data and should be unaffected by genotyping inaccuracies. Hence the high levels of balancing selection in SVs (DELs and INVs) relative to SNPs made evident in this analysis helps to validate earlier findings in this manuscript regarding the increased rates of heterozygosities in SV reads compared to SNPs. Because moderate frequency alleles are more likely to occur in the presence of balancing selection (Siewert & Voight 2017), these results suggest that the observed high SV Ho measures are legitimate and not entirely due to underlying genotyping inaccuracies from short-read SV calling, because they are supported by underlying patterns in the SNPs.

While we identified that balancing selection scores were higher in outlier SNPs than non-outlier SNPs, we did not find the difference to be significant (Supplementary Materials: Table S5), though the results of the outlier SNP PCA (Fig. 6c) compared to popgen-filtered SVs PCA (Fig. 4e) support this trend, as we see greater relative UK to AU spread in the former plot. We see much greater diversity in UK compared to AU individuals for outlier SNPs than with outlier SVs in the PCA plots (Fig. 6c&f); however, this observation may be due to the very low number of outlier SVs flagged, which provide poor resolution. Increasing sample sizes, particularly in the native range, would be likely to increase our ability to identify patterns of divergent and balancing selection across these populations.

We see the above discussed trends when the pseudo β^(1)^ scores were calculated using a slightly different approach that did not require scores to be averaged within each SV. However, under this second method, SV pseudo β^(1)^ scores are higher comparative to SNPs β^(1)^ scores (Supplementary Materials: Fig. S12, Table S6). It is likely that use of trimmed averages for each SV is the more appropriate approach, as the trimmed average for each SNP helps to reduce overcontribution of linked SNPs with high β^(1)^ scores inflating the pseudo β^(1)^ scores for the SVs.

Despite previous assumptions that balancing selection may be a rare phenomenon (Fijarczyk & Babik 2015), there is a growing body of literature reporting widespread instances of this phenomenon facilitating adaptation under different selection regimes (de Filippo *et al*. 2016; Huang *et al*. 2016; Koenig *et al*. 2019). A recent study of the invasive copepod *Eurytemora affinis* found that genetic variants with signatures of long-term balancing selection in heterogenous native ranges were more likely to undergo parallel adaptive evolution in response to shifted environmental salinity levels within invasive ranges (Stern & Lee 2020). With this growing appreciation of the role of balancing selection in facilitating rapid adaption comes a need to better understand how selection regime shifts (e.g. land use change, climate change) reduce genetic diversity via loss of variants previously under balancing selection. Examining selection regime shifts relative to previous balancing selection equilibriums will be an important avenue of investigation to understand how declining native populations, such as in the starling (Heldbjerg *et al*. 2016; Versluijs *et al*. 2016), may spiral further towards detrimental genetic diversity loss and collapse. Our results support the hypothesis that balancing selection plays a key role in maintain genetic variants and indicates that different types of genetic variants may experience different levels of balancing selection within a population. Expanding Australian and especially native range sampling in future studies, along with improved SV identification via complimentary long-read data, would enable us to further validate these findings.

## 5. Conclusion

The findings of our research demonstrate that even within recently diverged lineages or populations, there may be high amounts of SV variation. Further, patterns of SV genetic diversity do not necessarily reflect relative patterns in SNP diversity, either when considering patterns of diversity along the length of the organism’s chromosomes (owing to enrichment of SVs in sub telomeric repeat regions), or interpopulation diversity patterns (possibly a result of altered selection regimes or introduction history). Finally, we found that levels of balancing selection within the native range differed across SNP and SV groupings, suggesting that selection regimes may favour different optimum frequencies for different types of genetic variants. Overall, our results demonstrate that the processes that shape allelic diversity within populations is complex and demonstrates the need for further investigation of SVs across a range of taxa to better understand correlations between oft well studied SNP diversity and that of SVs.

## Notes

### Competing Interest Statement

The authors have declared no competing interest.

