## Supplementary Materials for "Contrasting patterns of single nucleotide polymorphisms and structural variations across multiple invasions"

**Table S1: Summary of WGS *Sturnus vulgaris* individuals** used for whole genome SNP and SV calling.

| IND. | Pop. Status | Country | Region | Location | AU subcluster | Total Reads |
|---|---|---|---|---|---|---|
| au01 | Invasive | Australia | NSW | Lemontree | East | 161,774,866 |
| au02 | Invasive | Australia | NSW | Lemontree | East | 138,481,596 |
| au03 | Invasive | Australia | NSW | Maitland | East | 121,949,997 |
| au04 | Invasive | Australia | NSW | Maitland | East | 121,934,492 |
| au05 | Invasive | Australia | SA | Meningie | South | 121,963,487 |
| au06 | Invasive | Australia | SA | Meningie | South | 130,652,648 |
| au07 | Invasive | Australia | VIC | Wonthaggi | South | 130,147,119 |
| au08 | Invasive | Australia | VIC | Wonthaggi | South | 159,551,759 |
| au09 | Invasive | Australia | WA | Munglinup | South | 168,989,017 |
| au10 | Invasive | Australia | NSW | Hay | East/South | 118,574,348 |
| au11 | Invasive | Australia | WA | Munglinup | South | 120,434,420 |
| au12 | Invasive | Australia | WA | Condingup | South | 115,005,268 |
| au13 | Invasive | Australia | NSW | Hay | East/South | 132,738,406 |
| au14 | Invasive | Australia | NSW | Dubbo | East | 110,957,661 |
| au15 | Invasive | Australia | SA | Meningie | South | 128,379,920 |
| au16 | Invasive | Australia | WA | Munglinup | South | 260,596,607 |
| au17 | Invasive | Australia | WA | Munglinup | South | 224,492,296 |
| au18 | Invasive | Australia | WA | Condingup | South | 211,097,422 |
| au19 | Invasive | Australia | WA | Condingup | South | 218,651,503 |
| au20 | Invasive | Australia | TAS | Hobart | South | 215,058,859 |
| au21 | Invasive | Australia | TAS | Hobart | South | 329,774,102 |
| au22 | Invasive | Australia | TAS | Hobart | South | 266,604,570 |
| au23 | Invasive | Australia | NSW | Dubbo | East | 198,731,521 |
| au24 | Invasive | Australia | NSW | Dubbo | East | 230,443,227 |
| au25 | Invasive | Australia | NSW | Hay | East/South | 211,675,237 |
| au26 | Invasive | Australia | NSW | Lemontree | East | 243,645,414 |
| au27 | Invasive | Australia | NSW | Maitland | East | 259,735,087 |
| au28 | Invasive | Australia | WA | Munglinup | South | 239,006,061 |
| au29 | Invasive | Australia | WA | Munglinup | South | 264,975,415 |
| au30 | Invasive | Australia | VIC | Wonthaggi | South | 235,789,158 |
| au31 | Invasive | Australia | WA | Munglinup | South | 390,701,211 |
| au32 | Invasive | Australia | WA | Munglinup | South | 630,512,935 |
| au33 | Invasive | Australia | WA | Munglinup | South | 503,636,307 |
| us01 | Invasive | USA | NY | New York | - | 170,518,757 |
| us02 | Invasive | USA | NY | New York | - | 127,905,364 |
| us03 | Invasive | USA | NY | New York | - | 104,086,567 |
| us04 | Invasive | USA | NY | New York | - | 126,123,487 |
| us05 | Invasive | USA | NY | New York | - | 103,209,440 |
| us06 | Invasive | USA | NY | New York | - | 140,354,835 |
| us07 | Invasive | USA | NY | New York | - | 150,480,727 |
| us08 | Invasive | USA | NY | New York | - | 132,168,818 |

| | | | | | | |
|---|---|---|---|---|---|---|
| uk01 | Native | England | UK | Newcastle upon Tyne | - | 122,402,512 |
| uk02 | Native | England | UK | Newcastle upon Tyne | - | 166,651,716 |
| uk03 | Native | England | UK | Newcastle upon Tyne | - | 160,423,348 |
| uk04 | Native | England | UK | Newcastle upon Tyne | - | 143,343,371 |
| uk05 | Native | England | UK | Newcastle upon Tyne | - | 134,177,188 |
| uk06 | Native | England | UK | Newcastle upon Tyne | - | 138,343,599 |
| uk07 | Native | England | UK | Newcastle upon Tyne | - | 154,792,665 |
| uk08 | Native | England | UK | Newcastle upon Tyne | - | 150,807,756 |

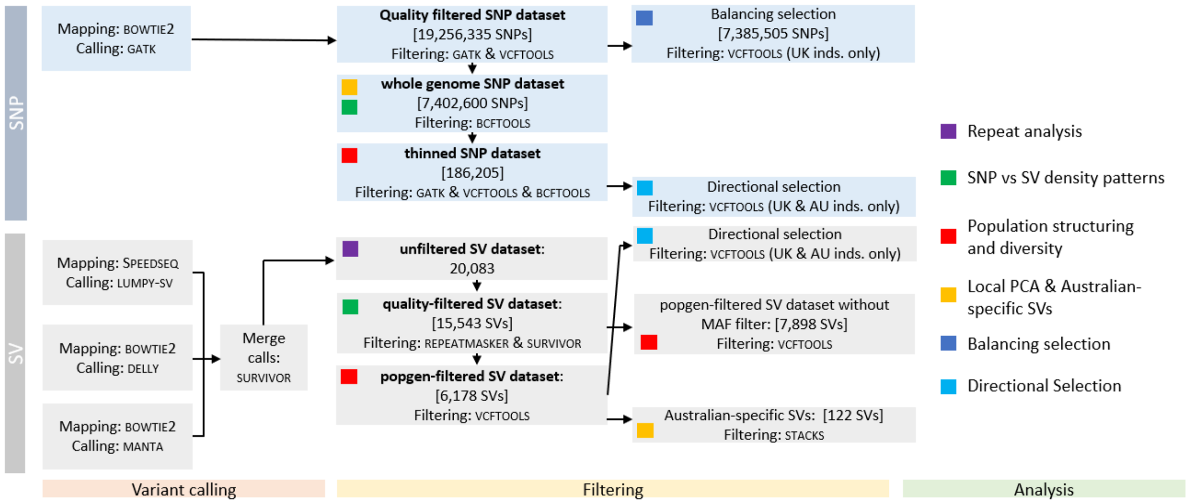


**Figure S1. A summary of variant calling and filtering used in various analysis within this manuscript.** Bold names of datasets in the 'filtering' section are named and referred to directly in the manuscript text. Additional datasets generated by extra filtering choices are also listed in this figure (see main body of manuscript for rationales and justifications).

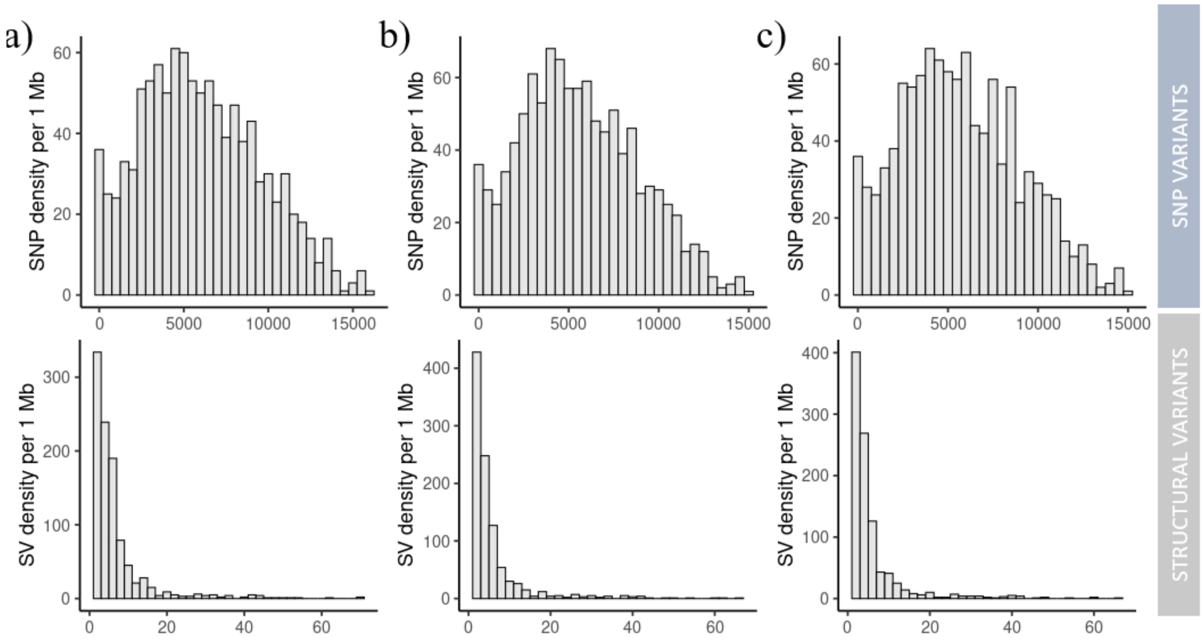


**Figure S2: Density histograms (1 Mb windows) of SNP and SV in *Sturnus vulgaris*** across the sampled continents, with panel a) invasive AU, panel b) invasive NA, and panel c) native range UK samples.

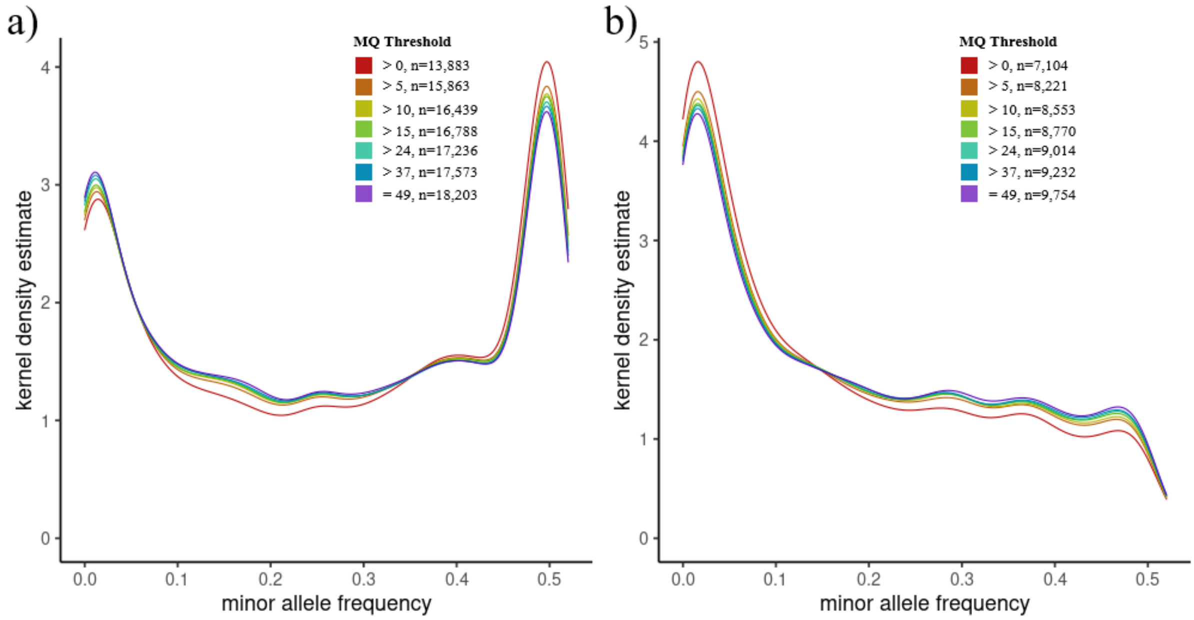


**Figure S3: Minor allele frequency kernel density estimated at different MQ thresholds for *Sturnus vulgaris* structural variants**, with panel a) unfiltered variants post MQ filtering, and panel b) variants filtered for 30 missingness across all individual post MQ filtering. Total number of variants plotted is indicated next to colour legends in each panel.

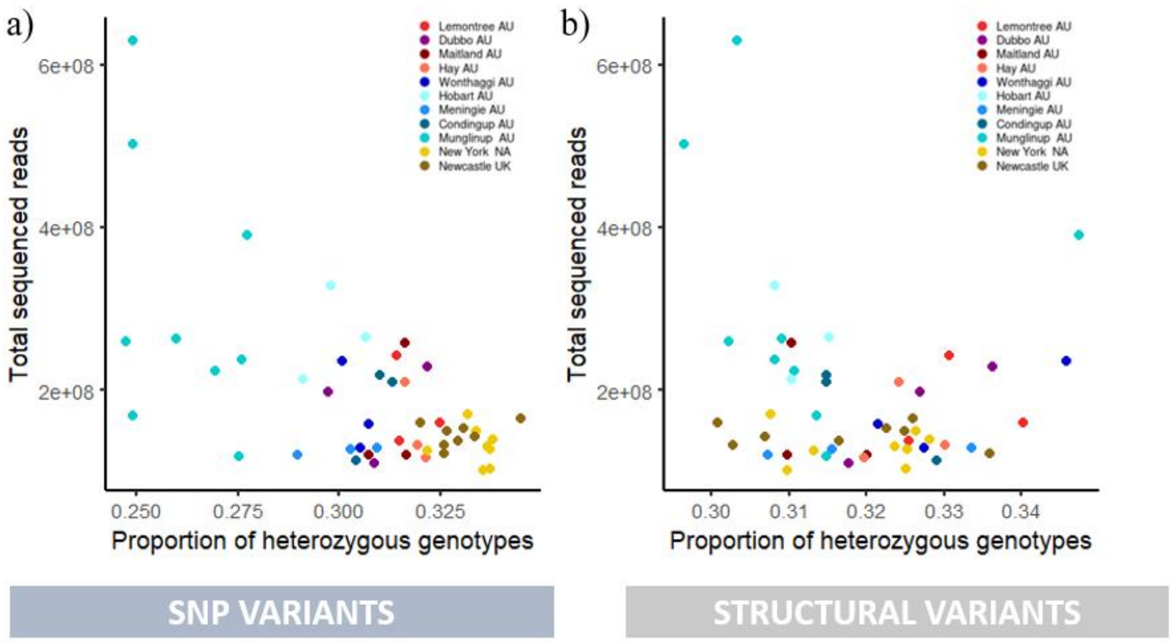


**Figure S4: Total sequenced read count verses heterozygous genotype proportion across Sturnus vulgaris** samples for SNPs and SV variants, coloured by sample site. We found no effect of read counts on individual heterozygosity rates using a linear mixed model, with population set as a random factor (SNP: β=-0.104, SE= 0.0613, z(49)= -1.692, SV: β=-0.190, SE=0.151, z(49)= -1.264).

**Table S2: Genetic diversity ($\pi$) stats for global *Sturnus vulgaris* samples sites**, corresponding to π values on Figure 3. Assessed using STACKS *population.*

| | **SNP** | | **SV** | |
|---|---|---|---|---|
| | **Pi** | **StdErr** | **Pi** | **StdErr** |
| **Lemontree** | 0.29 | 0.00056 | 0.32896 | 0.00327 |
| **Maitland** | 0.28996 | 0.00056 | 0.31696 | 0.00336 |
| **Dubbo** | 0.29013 | 0.00056 | 0.3249 | 0.00328 |
| **Hay** | 0.29099 | 0.00056 | 0.32541 | 0.0033 |
| **Hobart** | 0.27315 | 0.00057 | 0.309 | 0.00337 |
| **Wonthaggi** | 0.28063 | 0.00057 | 0.33228 | 0.00325 |
| **Meningie** | 0.28023 | 0.00057 | 0.31982 | 0.00331 |
| **Condingup** | 0.27881 | 0.00057 | 0.32116 | 0.00335 |
| **Munglinup** | 0.22468 | 0.00049 | 0.28997 | 0.00259 |
| **New York** | 0.30807 | 0.00042 | 0.29957 | 0.00252 |
| **Newcastle upon Tyne** | 0.31422 | 0.0004 | 0.30086 | 0.00248 |

**Table S3: Genetic diversity stats for *Sturnus vulgaris* sample sites and AU sample groupings,** corresponding to Table 1, with associated standard errors. Assessed using STACKS *population* (variant and fixed alllels used in calculations). SNP data set was the thinned SNP dataset (186,205 SNPs), and SV datasets were the popgen-filtered SV dataset (6,178 SVs), and an alternate version of this that had no minor allele frequency (MAF) filtering (7,898 SVs).

| | **SNP** (186,205 SNPs) | | | | | | **SV** (6,178 SVs) | | | | | |
|---|---|---|---|---|---|---|---|---|---|---|---|---|
| | Ho | StdErr | He | StdErr | Fis | StdErr | Ho | StdErr | He | StdErr | Fis | StdErr |
| Australia | 0.2716 | 0.0004 | 0.2777 | 0.0004 | 0.0290 | 0.0013 | 0.3274 | 0.0031 | 0.3123 | 0.0020 | -0.0041 | 0.0499 |
| Lemontree | 0.2935 | 0.0007 | 0.2416 | 0.0005 | -0.0062 | 0.0002 | 0.3396 | 0.0044 | 0.2663 | 0.0026 | -0.0143 | 0.0054 |
| Maitland | 0.2908 | 0.0007 | 0.2415 | 0.0005 | -0.0016 | 0.0002 | 0.3214 | 0.0043 | 0.2550 | 0.0026 | -0.0039 | 0.0058 |
| Dubbo | 0.2853 | 0.0007 | 0.2417 | 0.0005 | 0.0086 | 0.0002 | 0.3330 | 0.0044 | 0.2627 | 0.0026 | -0.0094 | 0.0057 |
| Hay | 0.2948 | 0.0007 | 0.2424 | 0.0005 | -0.0068 | 0.0002 | 0.3316 | 0.0043 | 0.2626 | 0.0026 | -0.0069 | 0.0058 |
| Wonthaggi | 0.2797 | 0.0007 | 0.2338 | 0.0005 | 0.0016 | 0.0002 | 0.3384 | 0.0043 | 0.2692 | 0.0026 | -0.0060 | 0.0054 |
| Hobart | 0.2736 | 0.0007 | 0.2276 | 0.0005 | -0.0007 | 0.0002 | 0.3195 | 0.0044 | 0.2494 | 0.0026 | -0.0134 | 0.0052 |
| Meningie | 0.2775 | 0.0007 | 0.2334 | 0.0005 | 0.0049 | 0.0002 | 0.3249 | 0.0043 | 0.2582 | 0.0026 | -0.0045 | 0.0058 |
| Condingup | 0.2838 | 0.0007 | 0.2322 | 0.0005 | -0.0090 | 0.0002 | 0.3278 | 0.0044 | 0.2588 | 0.0026 | -0.0071 | 0.0056 |
| Munglinup | 0.2363 | 0.0006 | 0.2122 | 0.0005 | -0.0272 | 0.0004 | 0.3221 | 0.0039 | 0.2713 | 0.0024 | -0.0544 | 0.0148 |
| Hofmeister et al. subset of 8 AU inds. | 0.2857 | 0.0005 | 0.2723 | 0.0004 | 0.0114 | 0.0005 | 0.3334 | 0.0036 | 0.2905 | 0.0022 | -0.0274 | 0.0150 |
| New York | 0.3108 | 0.0005 | 0.2888 | 0.0004 | -0.0060 | 0.0005 | 0.3279 | 0.0037 | 0.2768 | 0.0023 | -0.0466 | 0.0161 |
| Newcastle upon Tyne | 0.3152 | 0.0005 | 0.2946 | 0.0004 | -0.0020 | 0.0005 | 0.3297 | 0.0037 | 0.2787 | 0.0023 | -0.0476 | 0.0161 |

**Table S3 (CONT): Genetic diversity stats for *Sturnus vulgaris* sample sites and AU sample groupings,** corresponding to Table 1, with associated standard errors. Assessed using STACKS *population* (variant and fixed alllels used in calculations). SNP data set was the thinned SNP dataset (186,205 SNPs), and SV datasets were the popgen-filtered SV dataset (6,178 SVs), and an alternate version of this that had no minor allele frequency (MAF) filtering (7,898 SVs).

| | **SV** (7,898 SVs) | | | | | |
|---|---|---|---|---|---|---|
| | **Ho** | **StdErr** | **He** | **StdErr** | **Fis** | **StdErr** |
| Australia | 0.2587 | 0.0028 | 0.2469 | 0.0021 | -0.0018 | 0.0412 |
| Lemontree | 0.2688 | 0.0037 | 0.2108 | 0.0024 | -0.0112 | 0.0045 |
| Maitland | 0.2536 | 0.0037 | 0.2013 | 0.0024 | -0.0029 | 0.0049 |
| Dubbo | 0.2644 | 0.0037 | 0.2090 | 0.0023 | -0.0068 | 0.0048 |
| Hay | 0.2621 | 0.0037 | 0.2078 | 0.0024 | -0.0053 | 0.0048 |
| Wonthaggi | 0.2682 | 0.0037 | 0.2137 | 0.0024 | -0.0042 | 0.0045 |
| Hobart | 0.2526 | 0.0038 | 0.1972 | 0.0024 | -0.0106 | 0.0043 |
| Meningie | 0.2563 | 0.0037 | 0.2038 | 0.0023 | -0.0034 | 0.0048 |
| Condingup | 0.2589 | 0.0038 | 0.2044 | 0.0024 | -0.0057 | 0.0047 |
| Munglinup | 0.2533 | 0.0034 | 0.2135 | 0.0023 | -0.0422 | 0.0121 |
| Hofmeister et al. subset of 8 AU inds. | 0.2633 | 0.0032 | 0.2296 | 0.0022 | -0.0214 | 0.0126 |
| New York | 0.2601 | 0.0032 | 0.2199 | 0.0022 | -0.0365 | 0.0135 |
| Newcastle upon Tyne | 0.2619 | 0.0032 | 0.2217 | 0.0022 | -0.0374 | 0.0134 |

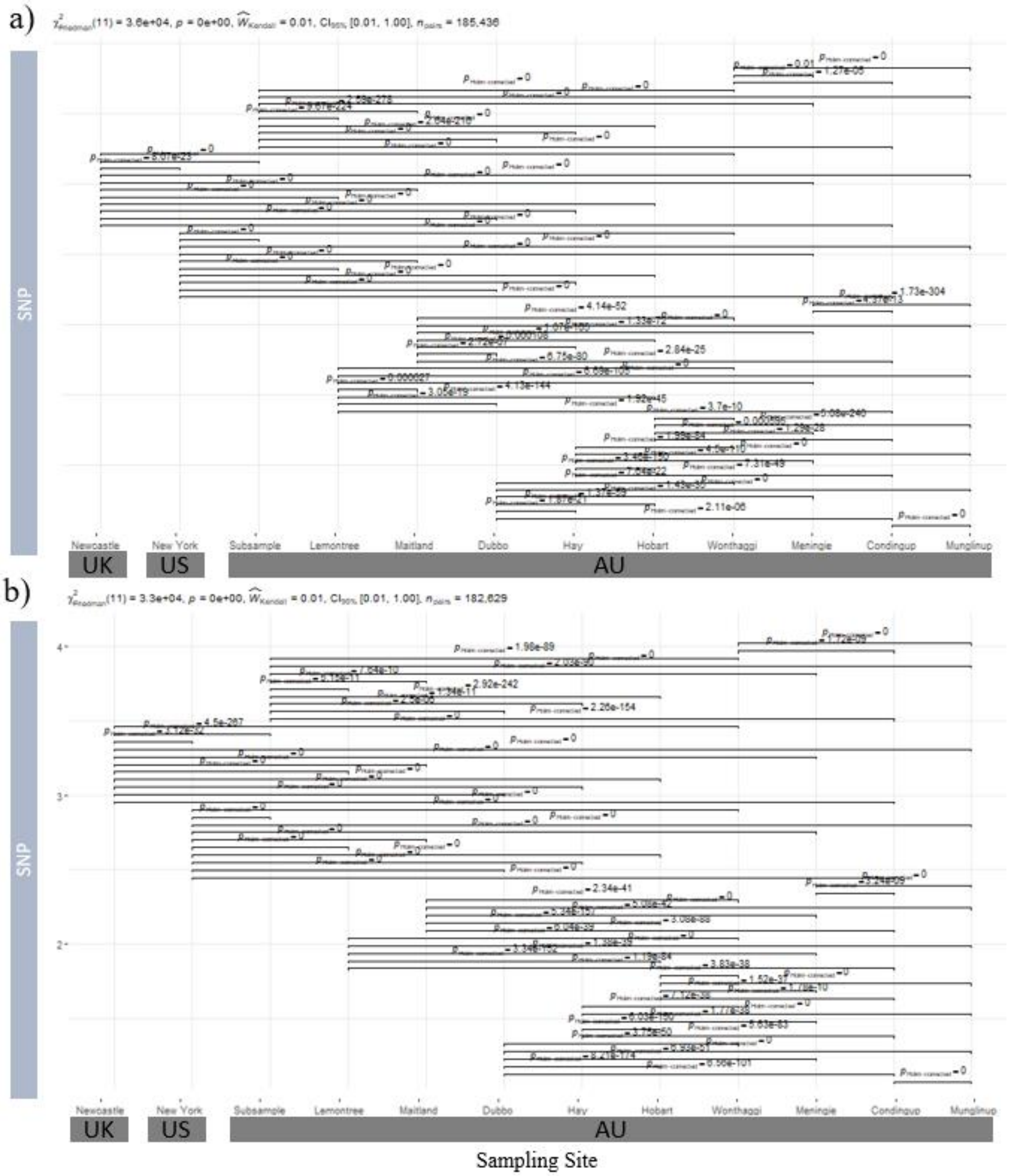


**Figure S5: Nonparametric within group comparison of genetic diversity measures across sample sites for popgen filtered SNPs (186,205) in *Sturnus vulgaris*,** with panel a) depicting Ho (observed heterozygosity) and panel b) He (expected heterozygosity). Pairwise testing via Durbin-Conover test, with only significant pairwise differences displayed.

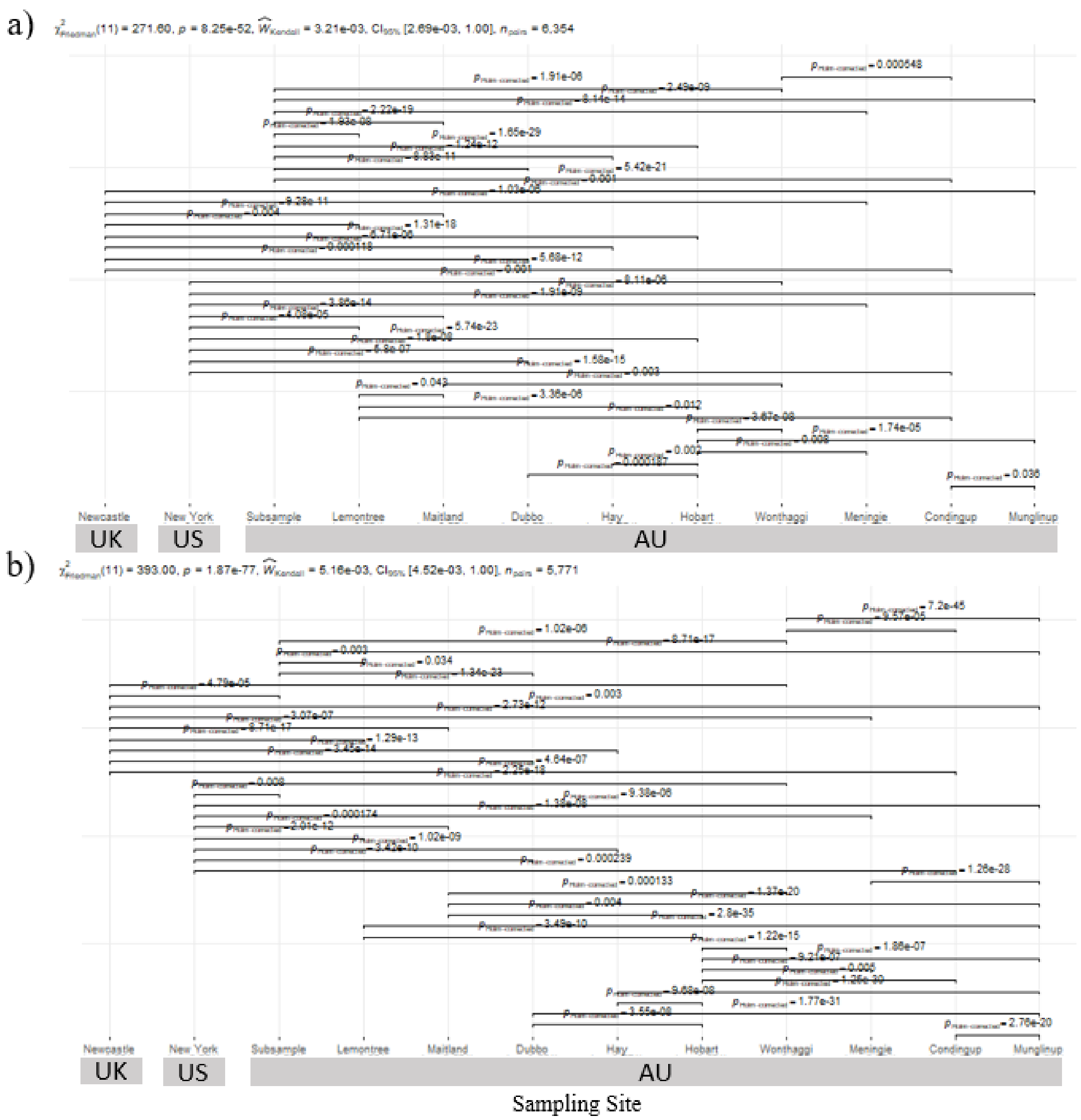


**Figure S6: Nonparametric within group comparison of genetic diversity measures across sample sites for popgen filtered structural variants (6,178) in *Sturnus vulgaris*,** with panel a) depicting Ho (observed heterozygosity) and panel b) He (expected heterozygosity). Pairwise testing via Durbin-Conover test, with only significant pairwise differences displayed.

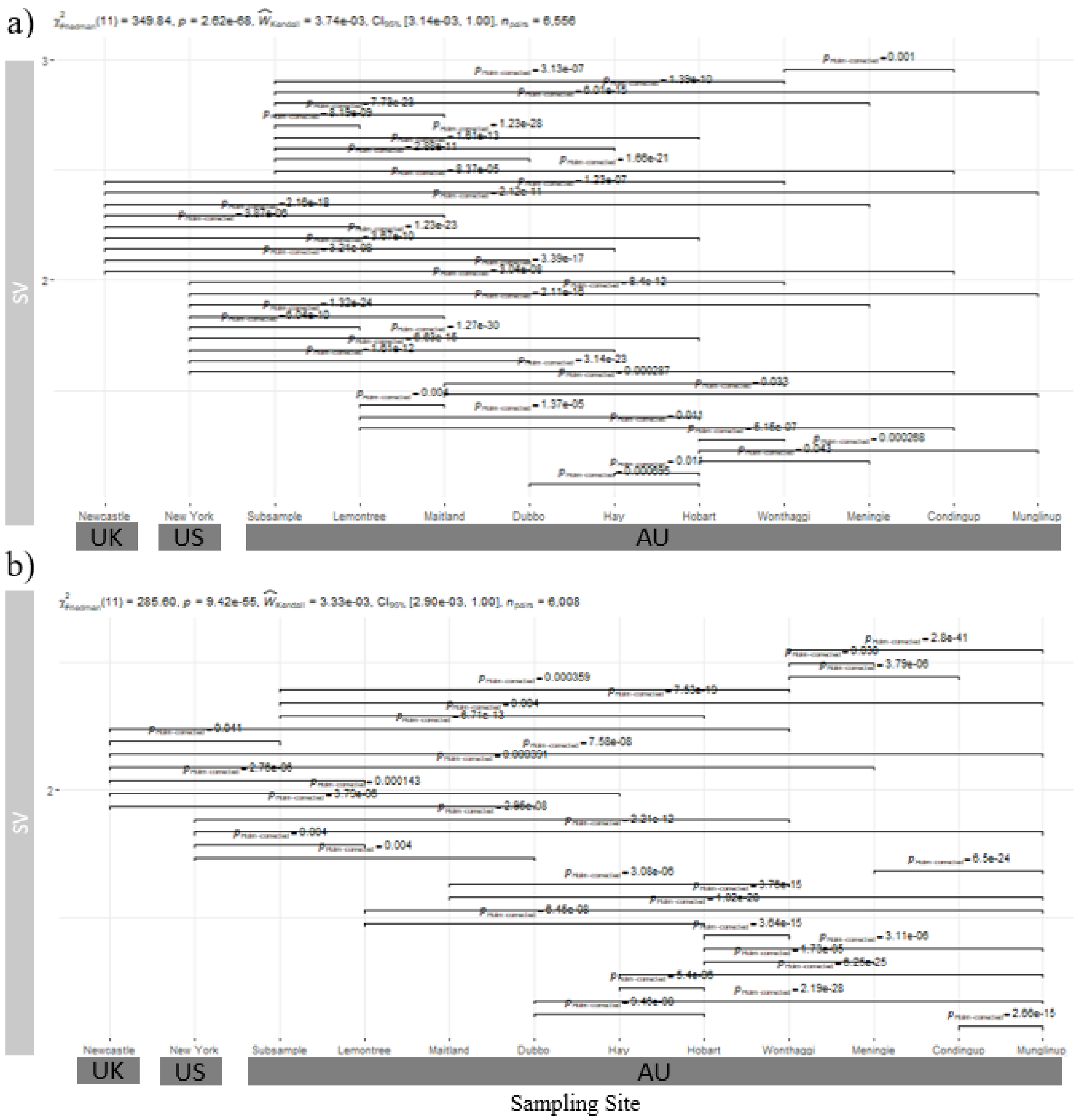


**Figure S7: Nonparametric within group comparison of genetic diversity measures across sample sites for non-maf filtered structural variants** (**7,898) in *Sturnus vulgaris*,** with panel a) depicting Ho (observed heterozygosity) and panel b) He (expected heterozygosity). Pairwise testing via Durbin-Conover test, with only significant pairwise differences displayed.

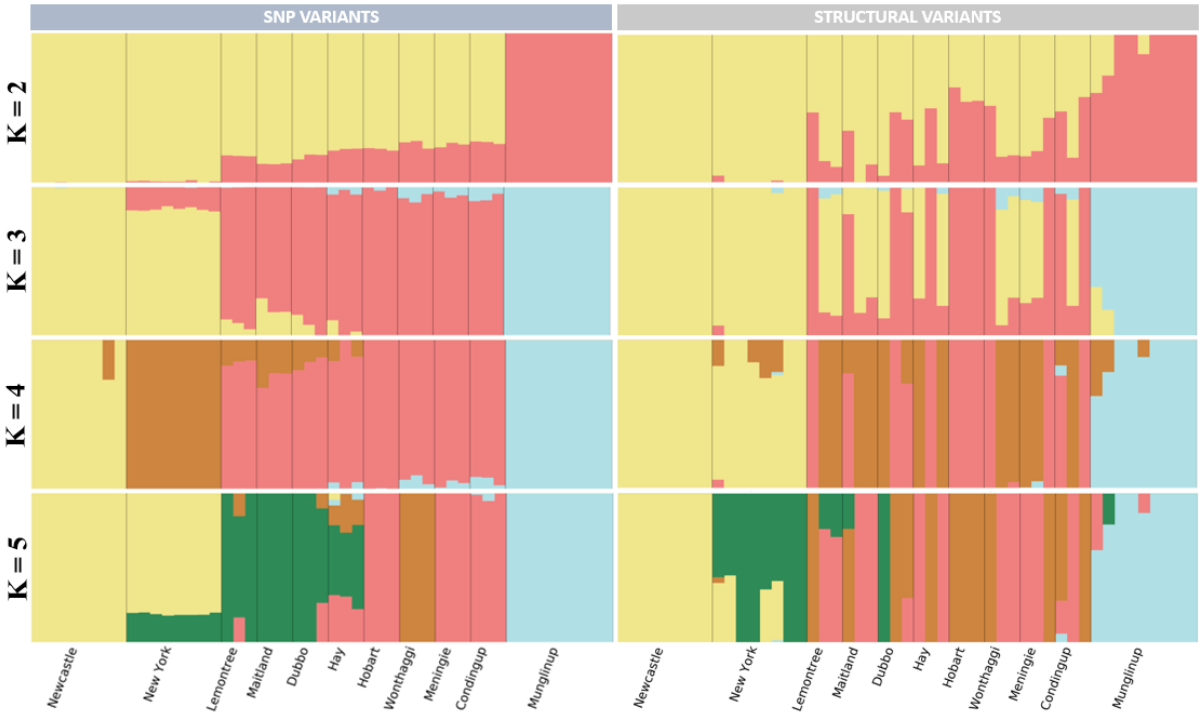


**Figure S8: Admixture plots SNP and SV in *Sturnus vulgaris*** across K values 2-5.

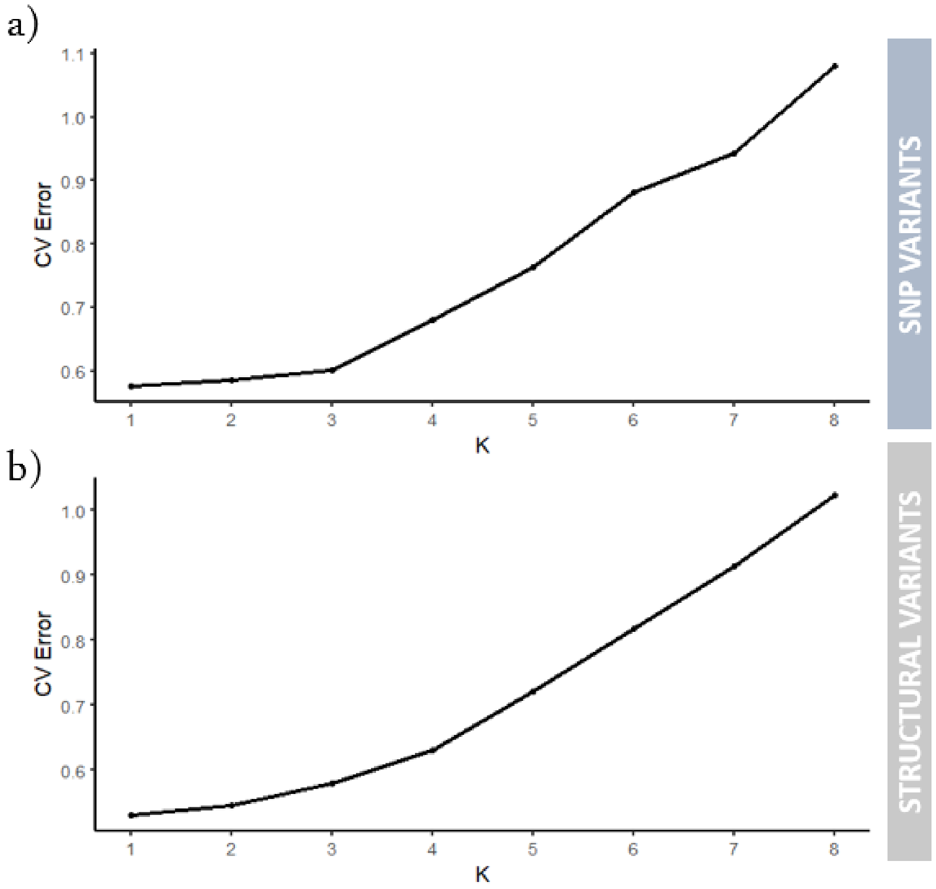


**Figure S9: Admixture cross-validation error plots SNP and SV in Sturnus vulgaris** across K values 1-8.

**Table S4: List of 1 kb plus common private SV alleles within Australia**, and the genes that they overlap with or exist within 1 kb of the ends of the structural variant.

| CHROM | POS | TYPE | LENGTH | GENES |
|---|---|---|---|---|
| chromosome7 | 33973495 | DEL | 24953 | LRP1B: Low-density lipoprotein receptor-related protein 1B |
| chromosome18 | 9879110 | DEL | 18056 | |
| chromosome19 | 28393 | DUP | 8970 | ASL: Argininosuccinate lyase<br>ASL2: Argininosuccinate lyase |
| chromosomeZ | 53804790 | DEL | 5028 | |
| chromosome17 | 2087800 | DEL | 5000 | RC3H2: Roquin-2<br>PHF19: PHD finger protein 19 |
| chromosome1 | 18226253 | DEL | 3714 | |
| chromosome3 | 15084714 | DEL | 3572 | |
| chromosome1A | 42402164 | DEL | 3000 | GPRC5A: Retinoic acid-induced protein 3 |
| chromosome2 | 115484306 | DEL | 2981 | Alkal1: ALK and LTK ligand 1 |
| chromosome1 | 18227437 | DUP | 2768 | |
| chromosome1 | 28848685 | DEL | 2209 | TAS2R40: Taste receptor type 2 member 40 |
| chromosome9 | 23431043 | DEL | 1803 | |
| chromosome20 | 15963693 | DEL | 1720 | |
| chromosome1 | 86600148 | DEL | 1650 | |
| chromosome3 | 57291865 | DEL | 1647 | |
| chromosome25 | 716229 | DEL | 1459 | Fcrla: Fc receptor-like A<br>CYP11B: Cytochrome P450 11B%2C mitochondrial<br>TAP2: Antigen peptide transporter 2 |
| chromosome8 | 6434687 | DEL | 1433 | |
| chromosome17 | 2919728 | DEL | 1277 | OLFM1: Noelin |
| chromosome1 | 46218927 | DEL | 1240 | |
| chromosome4 | 45401309 | DEL | 1171 | THEGL: Testicular haploid expressed gene protein-like |
| chromosome4 | 71146673 | DEL | 1156 | Ccdc158: Coiled-coil domain-containing protein 158 |
| chromosome1 | 75264936 | DEL | 1102 | MAOA: Amine oxidase [flavin-containing] A |
| chromosome2 | 123684082 | DEL | 1092 | ube2w: Ubiquitin-conjugating enzyme E2 W |
| chromosome4 | 3675969 | DEL | 1062 | |
| chromosome1A | 51123125 | DUP | 1060 | |
| chromosome2 | 79051315 | DEL | 1022 | IKZF1: DNA-binding protein Ikaros |

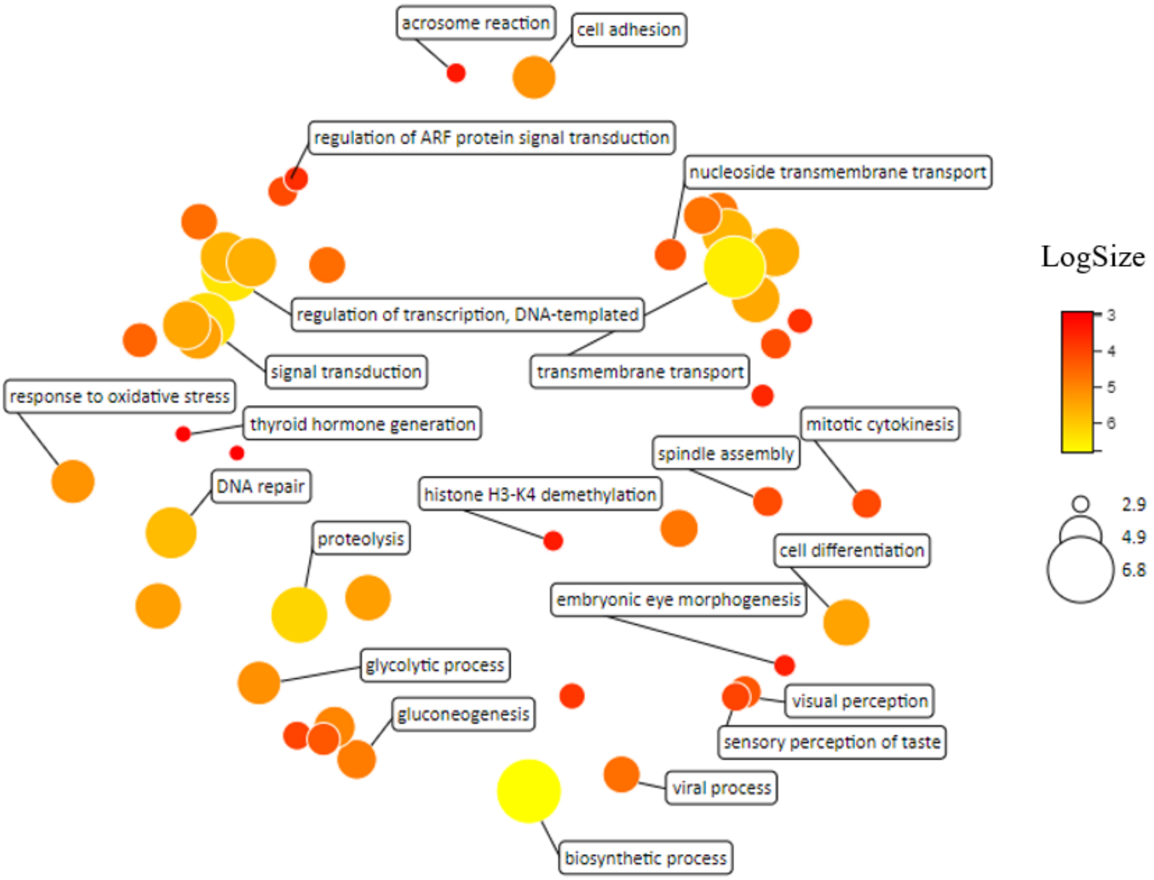


**Figure S10: Summary of gene ontology (GO) biological processes for common unique SVs in *Sturnus vulgaris*,** summarised using REVIGO. The scatterplot shows representative clusters of the GO analysis, with Log Size (indicated by circle diameter and colour) representing the frequency of GO term in the data set, plotted in semantic space with similar GO terms being placed close to one another.

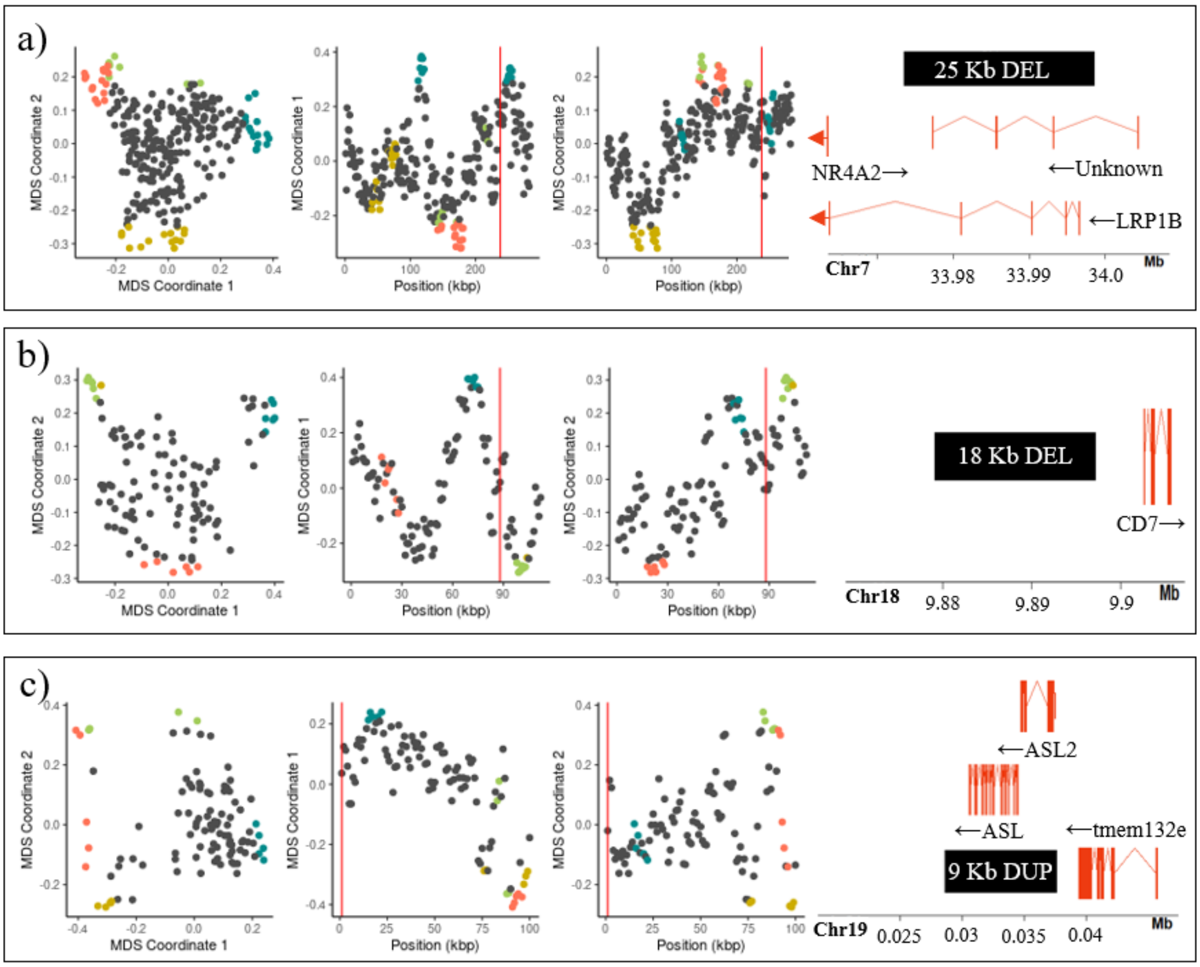


**Figure S11: Local PCA results for the genome wide single nucleotide polymorphism (SNP) dataset (1000 SNP windows), with the three largest Australian-specific structural variants (SVs) indicated as a red vertical line on top of local PCA plots, and their gene structure maps.** Panel (a) depicts a 25 kb DEL on chromosome 7. Panel (b) depicts an 18 kb DEL of chromosome 18. Panel (c) depicts a 9 kb DUP on chromosome 19. Within each panel, the first pane depicts the MDS visualisation of relationships between windows on the first two MDS axes, the second and third panes depicts the first two axes of MDS coordinates against the first SNP of each window with the red line denoting the position of the focal SV. Coloured SNP windows (green, blue, red) are consistent for plots within each panel and are used to help visualise tracks regions of extreme dissimilarity within each panel (obtained by colouring the top and bottom 5% SNP widows for MDS coordinate 1 and 2). Gene maps depict relative placement of the focal SVs (black) to genes (red) within 10 kb upstream and downstream, with black arrows indicating strand of the gene, solid red bars indicating gene exon, and connective lines between bars indicating introns within the gene (red arrows on gene maps indicate gene runs over the edge of the plotted window).

**Table S5: TukeyHSD's posthoc assessment of pairwise balancing selection differences between sample groups for SNPs and SVs under directional selection (outliers) and not under directional selection (nonoutliers) between AU and UK**, calculated in R using the *TukeyHSD()* function as a follow up to a significant *aov()* result ($F_{7, 113659}$ = 26.76, p-value < 0.0001). (Significant level indicated by: 0.05 > p-value ≥ 0.01 = *, 0.01 > p-value ≥ 0.001 = **, 0.001 > p-value = ***).

| **TukeyHSD** | Difference | 95% confidence interval | | Adjusted P-Value | Sig. |
|---|---|---|---|---|---|
| | | Lower | Upper | | |
| SNPnonoutlier-SNPoutlier | -0.524 | -1.354 | 0.306 | 0.542 | |
| SVoutlier-SNPoutlier | 0.181 | -1.756 | 2.118 | 1.000 | |
| SVnonoutliersDEL-SNPoutlier | -0.204 | -1.037 | 0.630 | 0.996 | |
| SVnonoutliersDUP-SNPoutlier | -0.216 | -1.155 | 0.723 | 0.997 | |
| SVnonoutliersINS-SNPoutlier | -0.116 | -3.920 | 3.688 | 1.000 | |
| SVnonoutliersTRA-SNPoutlier | -0.390 | -1.267 | 0.487 | 0.880 | |
| SVnonoutliersINV-SNPoutlier | 1.905 | -0.032 | 3.842 | 0.058 | |
| SVoutlier-SNPnonoutlier | 0.705 | -1.045 | 2.455 | 0.926 | |
| SVnonoutliersDEL-SNPnonoutlier | 0.320 | 0.243 | 0.397 | 0.000 | *** |
| SVnonoutliersDUP-SNPnonoutlier | 0.308 | -0.132 | 0.747 | 0.400 | |
| SVnonoutliersINS-SNPnonoutlier | 0.408 | -3.304 | 4.121 | 1.000 | |
| SVnonoutliersTRA-SNPnonoutlier | 0.134 | -0.150 | 0.417 | 0.843 | |
| SVnonoutliersINV-SNPnonoutlier | 2.429 | 0.679 | 4.179 | 0.001 | ** |
| SVnonoutliersDEL-SVoutlier | -0.384 | -2.136 | 1.367 | 0.998 | |
| SVnonoutliersDUP-SVoutlier | -0.397 | -2.201 | 1.407 | 0.998 | |
| SVnonoutliersINS-SVoutlier | -0.296 | -4.401 | 3.808 | 1.000 | |
| SVnonoutliersTRA-SVoutlier | -0.571 | -2.344 | 1.202 | 0.978 | |
| SVnonoutliersINV-SVoutlier | 1.724 | -0.751 | 4.199 | 0.407 | |
| SVnonoutliersDUP-SVnonoutliersDEL | -0.013 | -0.458 | 0.433 | 1.000 | |
| SVnonoutliersINS-SVnonoutliersDEL | 0.088 | -3.625 | 3.801 | 1.000 | |
| SVnonoutliersTRA-SVnonoutliersDEL | -0.187 | -0.479 | 0.106 | 0.530 | |
| SVnonoutliersINV-SVnonoutliersDEL | 2.108 | 0.357 | 3.860 | 0.006 | ** |
| SVnonoutliersINS-SVnonoutliersDUP | 0.101 | -3.638 | 3.839 | 1.000 | |
| SVnonoutliersTRA-SVnonoutliersDUP | -0.174 | -0.696 | 0.349 | 0.973 | |
| SVnonoutliersINV-SVnonoutliersDUP | 2.121 | 0.317 | 3.925 | 0.009 | ** |
| SVnonoutliersTRA-SVnonoutliersINS | -0.274 | -3.998 | 3.449 | 1.000 | |
| SVnonoutliersINV-SVnonoutliersINS | 2.020 | -2.084 | 6.125 | 0.812 | |
| SVnonoutliersINV-SVnonoutliersTRA | 2.295 | 0.522 | 4.068 | 0.002 | ** |

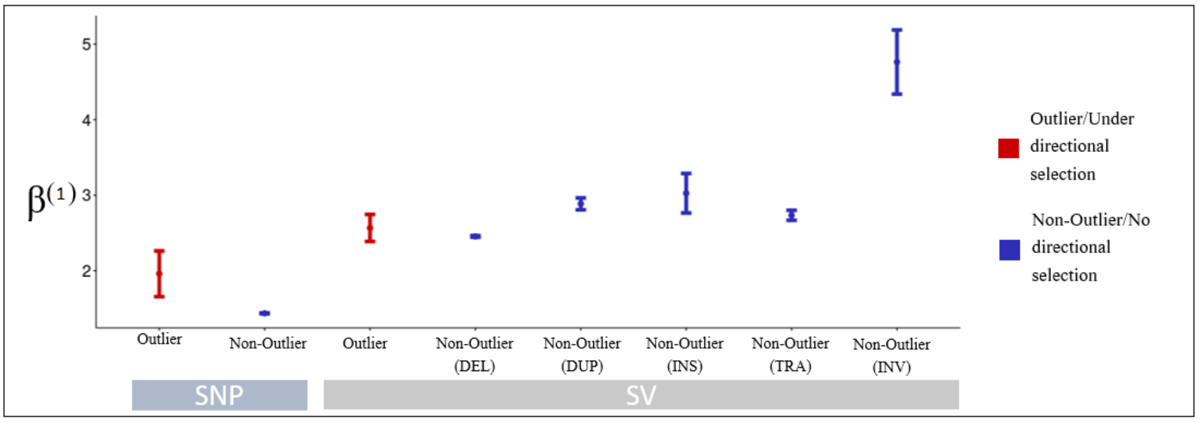


**Figure S12: Genetic variants under putative directional selection in *Sturnus vulgaris* across the native range and invasive Australian range and assessment of balancing selection,** with mean $\beta^{(1)}$ scores (balancing selection within the native UK range) for SNPs and SVs under directional selection (BAYESCAN outliers between AU and UK) and no directional selection (+/- standard error) in UK individuals, alongside the adjusted p-value of posthoc Tukey's pairwise tests of significance. SV pseudo $\beta^{(1)}$ scores calculated by averaging all overlapped SNP $\beta^{(1)}$ scores within each SV type group.

**Table S6: TukeyHSD's posthoc assessment of pairwise balancing selection differences between sample groups for SNPs and SVs under directional selection (outliers) and not under directional selection (nonoutliers) between AU and UK**, calculated in R using the *TukeyHSD()* function as a follow up to a significant *aov()* result ($F_{7,\ 158139} = 1036$, p-value < 0.0001). (Significant level indicated by: 0.05 > p-value ≥ 0.01 = *, 0.01 > p-value ≥ 0.001 = **, 0.001 > p-value = ***).

| **TukeyHSD** | Difference | 95% confidence interval | | Adjusted P-Value | Sig. |
|---|---|---|---|---|---|
| | | Lower | Upper | | |
| SNPnonoutlier-SNPoutlier | -0.524 | -1.662 | 0.614 | 0.860 | |
| SVoutlier-SNPoutlier | 1.162 | -0.235 | 2.559 | 0.187 | |
| SVnonoutliersDEL-SNPoutlier | 0.543 | -0.595 | 1.682 | 0.835 | |
| SVnonoutliersDUP-SNPoutlier | 0.274 | -0.881 | 1.430 | 0.996 | |
| SVnonoutliersINS-SNPoutlier | 1.112 | -0.662 | 2.886 | 0.551 | |
| SVnonoutliersTRA-SNPoutlier | 0.619 | -0.527 | 1.765 | 0.727 | |
| SVnonoutliersINV-SNPoutlier | 2.803 | 1.571 | 4.036 | 0.000 | *** |
| SVoutlier-SNPnonoutlier | 1.686 | 0.875 | 2.496 | 0.000 | *** |
| SVnonoutliersDEL-SNPnonoutlier | 1.067 | 1.027 | 1.108 | 0.000 | *** |
| SVnonoutliersDUP-SNPnonoutlier | 0.798 | 0.598 | 0.998 | 0.000 | *** |
| SVnonoutliersINS-SNPnonoutlier | 1.636 | 0.275 | 2.996 | 0.007 | ** |
| SVnonoutliersTRA-SNPnonoutlier | 1.143 | 1.010 | 1.276 | 0.000 | *** |
| SVnonoutliersINV-SNPnonoutlier | 3.327 | 2.853 | 3.801 | 0.000 | *** |
| SVnonoutliersDEL-SVoutlier | -0.618 | -1.429 | 0.192 | 0.287 | |
| SVnonoutliersDUP-SVoutlier | -0.888 | -1.722 | -0.054 | 0.028 | * |
| SVnonoutliersINS-SVoutlier | -0.050 | -1.633 | 1.533 | 1.000 | |
| SVnonoutliersTRA-SVoutlier | -0.543 | -1.363 | 0.278 | 0.478 | |
| SVnonoutliersINV-SVoutlier | 1.642 | 0.703 | 2.580 | 0.000 | *** |
| SVnonoutliersDUP-SVnonoutliersDEL | -0.269 | -0.471 | -0.067 | 0.001 | ** |
| SVnonoutliersINS-SVnonoutliersDEL | 0.568 | -0.792 | 1.929 | 0.911 | |
| SVnonoutliersTRA-SVnonoutliersDEL | 0.076 | -0.060 | 0.211 | 0.692 | |
| SVnonoutliersINV-SVnonoutliersDEL | 2.260 | 1.785 | 2.735 | 0.000 | *** |
| SVnonoutliersINS-SVnonoutliersDUP | 0.838 | -0.537 | 2.213 | 0.588 | |
| SVnonoutliersTRA-SVnonoutliersDUP | 0.345 | 0.107 | 0.583 | 0.000 | *** |
| SVnonoutliersINV-SVnonoutliersDUP | 2.529 | 2.016 | 3.043 | 0.000 | *** |
| SVnonoutliersTRA-SVnonoutliersINS | -0.493 | -1.860 | 0.874 | 0.958 | |
| SVnonoutliersINV-SVnonoutliersINS | 1.691 | 0.251 | 3.132 | 0.009 | ** |
| SVnonoutliersINV-SVnonoutliersTRA | 2.184 | 1.693 | 2.676 | 0.000 | *** |
